# Lipid kinases VPS34 and PIKfyve coordinate a phosphoinositide cascade to regulate Retriever-mediated recycling on endosomes

**DOI:** 10.1101/2021.05.25.445615

**Authors:** Sai Srinivas Panapakkam Giridharan, Guangming Luo, Pilar Rivero-Ríos, Noah Steinfeld, Helene Tronchere, Amika Singla, Ezra Burstein, Daniel D. Billadeau, Michael A. Sutton, Lois Weisman

**Affiliations:** Life Sciences Institute and Department of Cellular and Developmental Biology, University of Michigan, Ann Arbor, MI; INSERM U1048 I2MC, Toulouse, France and Université Paul Sabatier, Toulouse, France; Department of Internal Medicine, and Department of Molecular Biology, University of Texas Southwestern Medical Center, Dallas, TX; Division of Oncology Research and Schulze Center for Novel Therapeutics, Mayo Clinic, Rochester, MN; Neuroscience Graduate Program, Molecular and Behavioral Neuroscience Institute, Molecular and Integrative Physiology, University of Michigan, Ann Arbor, MI

**Author notes:** equal contributors (Sai Srinivas Panapakkam Giridharan, Guangming Luo).

**Keywords:** phosphatidylinositol 3, 5-bisphosphate, PtdIns(3,5)P_2_, PtdIns5P, Retriever, SNX17, VPS35L/C16orf62, VPS26C/DSCR3, CCC, COMMD, PIKfyve, VPS34

## Abstract

Cell-surface receptors control how cells respond to their environment. Many cell-surface receptors recycle from endosomes to the plasma membrane via a recently discovered pathway, which includes sorting-nexin SNX17, Retriever, WASH and CCC complexes. Here we discover that PIKfyve and its upstream PI3-kinase VPS34 positively regulate this pathway. VPS34 produces PI3P, which is the substrate for PIKfyve to generate PI3,5P_2_. We show that PIKfyve controls recycling of cargoes including integrins, receptors that control cell migration. Furthermore, endogenous PIKfyve colocalizes with SNX17, Retriever, WASH and CCC complexes on endosomes. Importantly, PIKfyve inhibition causes a loss of Retriever and CCC from endosomes, and mutation of the lipid binding site on a CCC subunit impairs its endosomal localization and delays integrin recycling. In addition, we show that recruitment of SNX17 is an early step and requires VPS34. These discoveries suggest that VPS34 and PIKfyve coordinate an ordered pathway to regulate recycling from endosomes and suggest how PIKfyve functions in cell migration.

## Introduction

The functions of many cell-surface receptors are controlled in part via the regulation of their exposure to the cell surface. Receptors are removed from the cell surface via regulated endocytosis, and then are either returned via recycling pathways or sent to lysosomes for degradation (Cullen and Steinberg, 2018; Grant and Donaldson, 2009; Naslavsky and Caplan, 2018). Multiple types of receptors are regulated via endocytosis and regulated recycling. These include G-protein-coupled receptors (GPCRs), post-synaptic receptors for neurotransmitters, nutrient transporters, and cell adhesion proteins. Thus, gaining molecular insight into the regulation of endocytosis and mechanisms for receptor recycling is key to understanding the control of multiple physiological processes.

A major recycling pathway was recently discovered that is regulated in part by the Retriever complex. The Retriever complex is composed of three proteins, VPS29, which is also in the retromer complex, and two unique subunits, VPS35L (C16orf62) and VPS26C (DSCR3). The Retriever complex acts with the sorting nexin, SNX17, the WASH complex, and the CCC complex which includes CCDC22 (coiled-coil domain containing 22)–CCDC93 (coiled-coil domain containing 93), and any one of ten COMMD (copper metabolism MURR1 domain)-containing proteins (Chen et al., 2019; McNally and Cullen, 2018; Simonetti and Cullen, 2019; Wang et al., 2018) to regulate trafficking of cargoes and membranes from early endosomes back to the cell surface. Proteomic studies revealed that this pathway traffics approximately half of the cell surface proteins in HeLa cells (McNally et al., 2017), while most of the remaining cell surface proteins are trafficked by the retromer pathway (Steinberg et al., 2013). Thus the Retriever pathway is a common route for protein recycling to the plasma membrane.

The best characterized cargoes of the SNX17-Retriever-CCC-WASH pathway are the integrins, which are transmembrane proteins that control cell migration via regulation of focal adhesion complexes (Wozniak et al., 2004). Integrins that are exposed on the cell surface connect the intracellular actin network to the extracellular matrix (Vicente-Manzanares et al., 2009). Integrin levels at the cell surface are controlled both by their endocytosis into endosomes, and their subsequent recycling back to the plasma membrane (Moreno-Layseca et al., 2019). Thus, understanding how integrin recycling is controlled is of great interest.

Control of the SNX17-Retriever-CCC-WASH pathway likely occurs in part via SNX17 recognition of cargoes as well as SNX17 association with the other proteins of the transport machinery. SNX17 specifically binds β1-integrin as well as other cargo receptors via direct interaction with an NPxY motif on the cytoplasmic side of the cargo protein (Bottcher et al., 2012; Chandra et al., 2019; Jia et al., 2014; McNally et al., 2017; Steinberg et al., 2012). In addition, SNX17 binds PI3P in vitro (Chandra et al., 2019). The WASH complex is also likely recruited to endosomes in part by binding to PI3P via the FAM21 subunit (Jia et al., 2010).

PI3P is one of seven phosphorylated phosphatidylinositol (PPI), which are low abundance signaling lipids that play multiple essential roles (De Craene et al., 2017; Schink et al., 2016). The seven PPI species are defined by their phosphorylation status at positions 3, 4, 5 of the inositol ring. The synthesis and turnover of PPIs are spatially and temporally regulated by several lipid kinases and phosphatases. PPI lipids act via the recruitment and control of their downstream effector proteins and orchestrate various cellular events including cell signaling, cytoskeletal organization, membrane trafficking, cell migration, and cell division (Dickson and Hille, 2019).

Some of the cellular PI3P serves as a substrate for PIKfyve, the lipid kinase that generates phosphatidylinositol 3,5-bisphosphate (PI3,5P_2_) (Hasegawa et al., 2017; Ho et al., 2012; McCartney et al., 2014a; Shisheva, 2012). PIKfyve also serves as the primary source for cellular pools of phosphatidylinositol 5-phosphate (PI5P), likely via the action of lipid phosphatases on PI3,5P_2_ (Zolov et al., 2012) and possibly by direct synthesis (Shisheva, 2012). PIKfyve exists in a protein complex which includes the lipid phosphatase, Fig4 and scaffold protein, Vac14. Both Fig4 and Vac14 positively regulate PIKfyve kinase activity (de Araujo et al., 2020; Dove et al., 2009; Ho et al., 2012; McCartney et al., 2014a; Shisheva, 2012; Strunk et al., 2020). Loss of PIKfyve or its positive regulators causes defects in lysosomal homeostasis and results in enlargement of endosomes and lysosomes in a variety of cell types (de Araujo et al., 2020; Dove et al., 2009; Ho et al., 2012; McCartney et al., 2014a; Shisheva, 2012).

The PIKfyve pathway is critical for normal function of multiple organs and tissues. Fig4 or Vac14 knock out mice die perinatally and exhibit profound neurodegeneration (Chow et al., 2007; Zhang et al., 2007). Moreover, mutations in *FIG4* and *VAC14* have been linked to neurological diseases including Charcot-Marie-Tooth syndrome and amyotrophic lateral sclerosis. Moreover, homozygous null mutations in human *FIG4* lead to infantile death and impairment of multiple organs (Lenk et al., 2016; McCartney et al., 2014a). Similarly, a PIKfyve hypomorphic mouse mutant dies perinatally and in addition to neurodegeneration has defects in multiple organs including the heart, lung and kidneys (Zolov et al., 2012). A whole body knock-out of PIKfyve results in very early lethality (Ikonomov et al., 2011; Takasuga et al., 2013). Together these studies indicate that PIKfyve is required for multiple physiological pathways and functions in multiple cell-types.

Recent studies indicate that PIKfyve plays roles in cell migration and invasion (Cinato et al., 2021; Dayam et al., 2017; Dupuis-Coronas et al., 2011; Oppelt et al., 2014; Oppelt et al., 2013; Shi and Wang, 2018). However, a full understanding of the molecular mechanisms whereby PIKfyve regulates cell migration remain to be established. Here we show that PIKfyve activity is required for cell migration. We discover that this occurs in part through regulation of the surface levels of β1-integrin via PIKfyve control of the SNX17-Retriever-CCC-WASH pathway.

## Results

### PIKfyve positively regulates cell migration

Recent studies using siRNA silencing as well as inhibition of PIKfyve indicate that PIKfyve plays a role in cell migration (Cinato et al., 2021; Oppelt et al., 2014; Oppelt et al., 2013; Shi and Wang, 2018). To further investigate, we performed wound-healing assays in the presence of two chemically distinct PIKfyve inhibitors, YM201636 and apilimod. In cultured cells, compared to vehicle control, PIKfyve inhibition with apilimod and YM201636 substantially delayed wound-healing by 45% and 40%, respectively (Figure 1A-B). Importantly, inhibition of PIKfyve did not significantly affect cell viability or proliferation (Figure S1), indicating that the delay in wound healing reflects a decrease in the ability of the cells to migrate.

**Figure 1.**
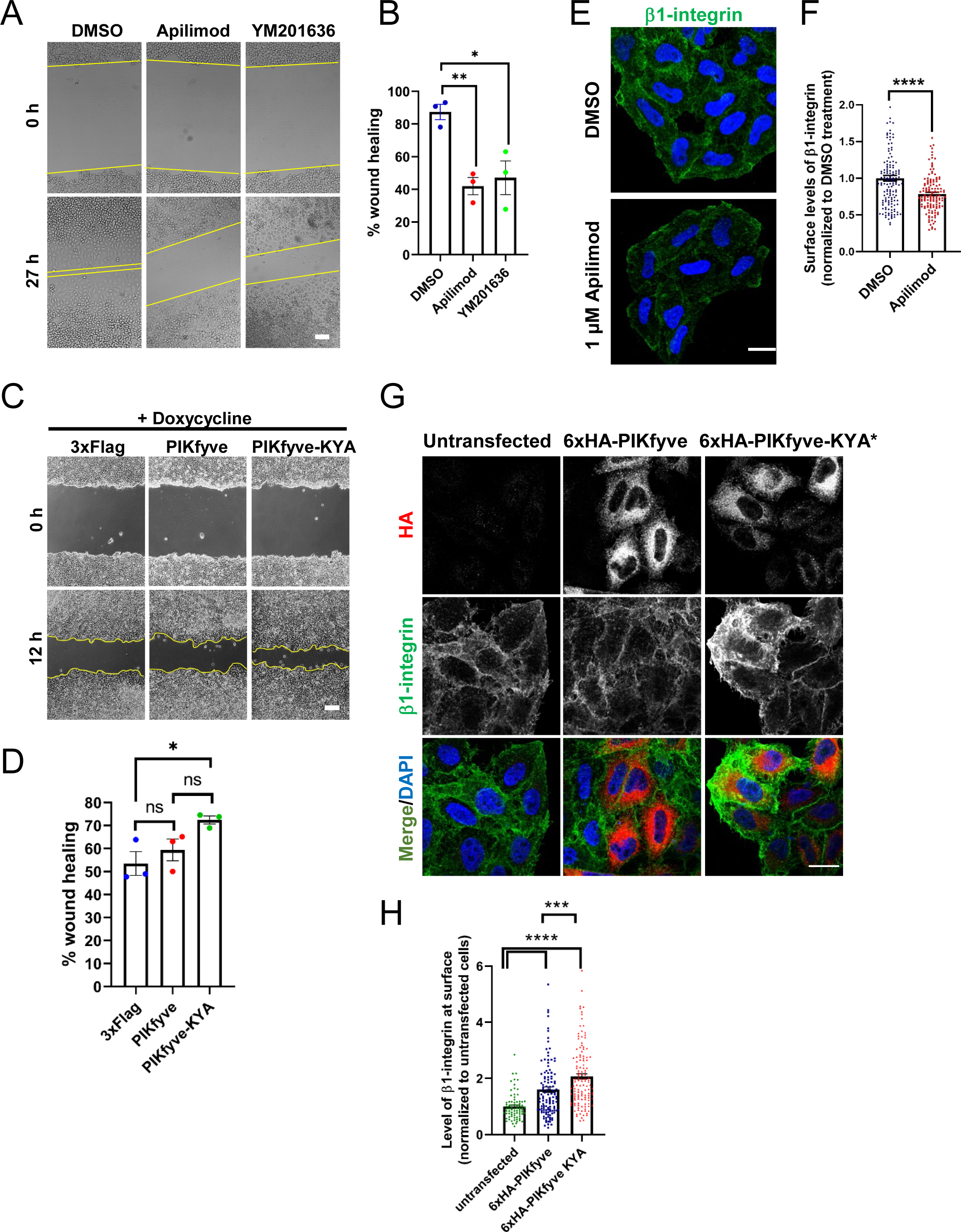
PIKfyve positively regulates cell migration in part via regulation of cell surface levels of β1-integrin. A-B Inhibition of PIKfyve delays cell migration. (A) Wound healing assays were performed on HeLa cells in the presence of either DMSO, 1 µM apilimod or 0.8 µM YM201636 for 27 h. (B) Percentage of wound closure was quantified. Bar-100 µm. C-D. Increasing PIKfyve activity promotes cell migration. (C) Wound healing assays were performed in Flp-in HEK293T cells stably expressing doxycycline-inducible wild-type PIKfyve or hyperactive PIKfyve-KYA in the presence of 100 ng/ml doxycycline for 12 h. (D) Percentage of wound area closure was quantified. Bar-100 µm. E-F. Inhibition of PIKfyve decreases the surface levels of β1-integrin. (E) HeLa cells treated with DMSO or 1 µM apilimod for 1 h were incubated with antibodies to label surface β1-integrin for 1 h at 4^0^C and fixed at 4^0^C. (F) Intensity of β1-integrin per cell was quantified and normalized to the average intensity of the DMSO treatment for that particular experiment. Bar-20 µm. G-H. Increasing PIKfyve activity elevates the surface levels of β1-integrin. (G) HeLa cells that were either untransfected or transiently transfected with 6xHA-PIKfyve or 6xHA-PIKfyve-KYA were incubated with antibodies to label surface β1-integrin for 1 h at 4^0^C. Cells were fixed, permeabilized, immunostained with an anti-HA antibody and corresponding Alexa-Fluor conjugated secondary antibodies. (H) Intensity of β1-integrin per cell was quantified and the values were normalized to the average intensity of untransfected cells for each experiment. Bar-20 µm. Data presented as mean ± SE. Statistical significance from three independent experiments were determined using unpaired two-tailed Student’s T-test (F) or one-way ANOVA and Dunnett’s (B) or Tukey (D,H) post hoc tests. Yellow lines indicate the migration front. * P<0.05, ** P<0.01, *** P<0.005, **** P<0.001 and ns, not significant.

We also observed a role for PIKfyve in cell migration in primary cells. Primary neonatal cardiac fibroblasts were analyzed in wound healing assays. Inhibition of PIKfyve with YM201636 impaired wound closure by 21% as compared to DMSO-treated cardiac fibroblasts (Figure S2A-B). Similar results were observed with apilimod (Cinato et al., 2021). In addition to chemical inhibition, we sought genetic evidence, and performed wound-healing experiments with primary fibroblasts derived from the hypomorphic PIKfyve^β-geo/β-geo^ mouse mutant, which expresses about 10% of the wild-type levels of PIKfyve and half the levels of PI3,5P_2_ and PI5P (Zolov et al., 2012). PIKfyve^β-geo/β-geo^ fibroblasts showed strong defects in cell migration and exhibited reduced wound healing by 30% compared to wild-type fibroblasts (Figure S2C-D).

Cell spreading is important for cell migration. Therefore, we assessed the ability of newly seeded cells to spread on plastic dishes coated with fibronectin, a component of the extracellular matrix. We found that inhibition of PIKfyve reduced cell spreading by 22% as compared to DMSO-treated control cells (Figure S2E-F).

The above studies, as well as previous studies, relied solely on lowering PIKfyve activity. To further test whether PIKfyve has a direct role in cell migration, we took the converse approach and tested whether elevation of PIKfyve activity promotes cell migration. We identified a hyperactive allele, PIKfyve-KYA, which elevates PI3,5P_2_ and PI5P 4-fold and 2-fold, respectively (McCartney et al., 2014a). Inducible expression of PIKfyve-KYA and wild-type PIKfyve was achieved via engineered Flp-In HEK293T cell lines. Induction of PIKfyve-KYA caused a significant increase in wound healing, by approximately 20%, whereas induction of wild-type PIKfyve resulted in a similar degree of cell migration to that observed in control cells (Figure 1C-D). The wound healing of these three cell lines was not different in the absence of doxycycline induction (Figure S2G-H). Together, these results suggest that PIKfyve activity directly impacts cell migration.

### PIKfyve activity plays a role in controlling surface levels of β1-integrin

That PIKfyve plays a direct role in cell migration, provided an opportunity to determine specific pathways that are directly regulated by PIKfyve. Integrins play a crucial role in cell migration. Thus, we tested whether PIKfyve activity correlates with integrin localization. Integrins are heterodimers of α and β chains, and β1-integrin is the most commonly found integrin β subunit (Moreno-Layseca et al., 2019). We therefore tested whether PIKfyve activity plays a role in the levels of β1-integrin at the cell surface. Notably, we found that when compared with DMSO-treated cells, apilimod treatment for 1 h, resulted in 21% less surface exposed β1-integrin (Figure 1E-F).

Conversely, we tested whether an increase in PIKfyve activity increases the surface levels of β1-integrin. We transiently expressed wild-type PIKfyve and PIKfyve-KYA in HeLa cells, and observed a 60% and 107% increase respectively, in β1-integrin on the cell surface as compared to untransfected control cells (Figure 1G-H). These findings suggest that PIKfyve activity regulates cell migration in part by regulating surface levels of β1-integrin.

### Inhibition of PIKfyve results in the accumulation of β1-integrin within internal compartments

That the surface levels of β1-integrin decrease following just 1 hour of PIKfyve inhibition, suggests that PIKfyve has a role in the dynamic regulation of integrin levels at the cell surface. To test this further, we determined the fate of surface β1-integrin after acute inhibition of PIKfyve for 15 or 30 min. We labeled HeLa cells with an antibody against an extracellular epitope of β1-integrin and assessed the levels of surface exposed and internalized β1-integrin in the presence of either DMSO or apilimod (Figure 2). Using methods to solely label surface integrin, we found that many of the cells were permeabilized during incubation on ice and subsequent fixation. Thus, we used an indirect approach. We fixed and permeabilized the cells, performed immunofluorescence localization of total β1-integrin. To estimate surface β1-integrin, we measured the amount within 0.8 µm of the cell border. In untreated cells, the percent of surface β1-integrin at the cell border was 25%. Following 30 min of incubation in DMSO, the amount of labeled integrin at the cell border exhibited a modest decrease to 21% of total integrin. In contrast, inhibition of PIKfyve for 30 min, caused a much larger decrease in the amount of integrin at the cell border to 13.7%. There was also a trend in the decrease in β1-integrin at the cell border following 15 min of apilimod treatment, but this change was not statistically significant (Figure 2K).

**Figure 2:**
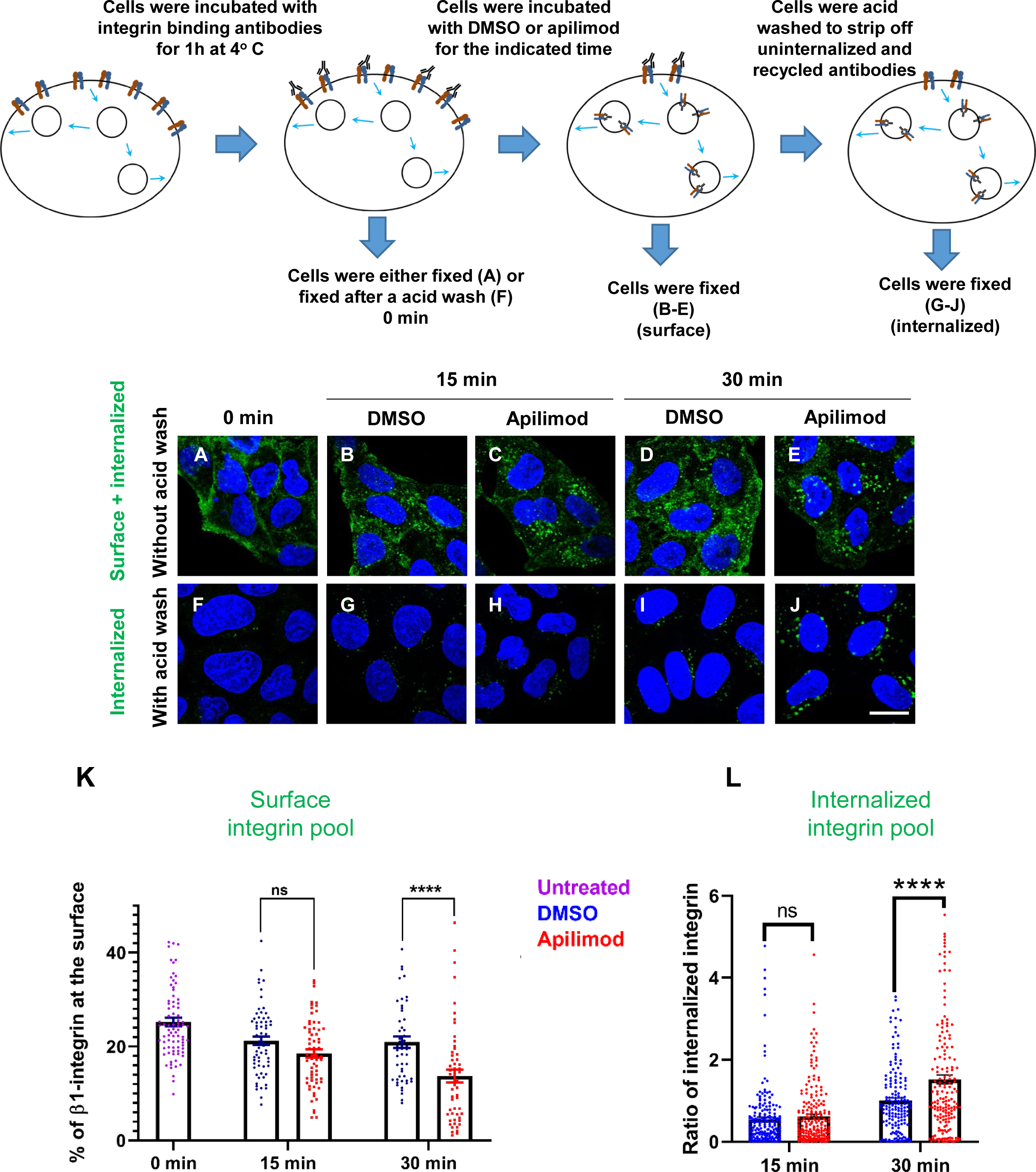
Inhibition of PIKfyve causes a rapid loss of β1-integrin from the cell surface and a concomitant accumulation of β1-integrin in internal compartments. A-J. HeLa cells were incubated with antibodies to label surface β1-integrin for 1 h at 4°C. Cells were either fixed (A), fixed after an acid wash (F), or incubated with media containing DMSO or 1 µM apilimod at 37°C for the indicated times. Following incubation, cells were fixed (B-E) or fixed after an acid wash (G-J). Fixed cells were permeabilized and immunostained with Alexa-Fluor-488 conjugated anti-mouse secondary antibodies. Flow diagram (top) outlines the experiment. K. The surface levels of β1-integrin were inferred from the intensity of β1-integrin within 0.8 micron from the cell border. Surface β1-integrin (for images A-E) is reported as the percentage of the total labeled β1-integrin. L. Internalized β1-integrin was quantified from cells treated as described in (G-J). β1-integrin intensity was normalized to the average intensity of cells treated with DMSO for 30 min for each experiment. Data presented mean ± SE. Statistical significance from three independent experiments was analyzed using two-way ANOVA and Sidak’s multiple comparisons tests. (K-L). *** P<0.005, **** P<0.001 and ns, not significant. Bar-10 µm.

In parallel, we measured the amount of labeled surface integrin that was internalized. We used a brief acid wash to remove surface β1-integrin bound antibodies and quantitated the protected, internalized β1-integrin bound to antibody (Figure 2F-J). Consistent with the observed changes in surface levels of integrin, we found that after 30 min of apilimod treatment, there was a significant increase of 0.52-fold more internalized β1-integrin as compared to DMSO-treated cells (Figure 2L). There was also a trend toward an increase in the internalized pool of β1-integrin after a 15 min treatment, although this change was not statistically significant. These results indicate that PIKfyve has a role in β1-integrin trafficking.

### Endogenous PIKfyve localizes to multiple endosomal compartments

To identify intracellular compartment(s) that act in PIKfyve-mediated regulation of β1-integrin recycling, we determined the localization of endogenous PIKfyve. To avoid overexpression-based artifacts, we generated a CRISPR-Cas9-engineered HEK293 cell line that expresses 3xHA-PIKfyve from the endogenous PIKfyve locus. We assessed the colocalization of endogenous 3xHA-PIKfyve with several endo-lysosomal markers including EEA1 (early endosomes), RAB7 (late endosomes), RAB11 (recycling endosomes), LAMP1 and LAMP2 (late endosomes/lysosomes) and VPS35 (retromer complex). We observed the best colocalization with VPS35 (Figure 3). VPS35 associates with endosomes that are active in retromer-dependent transport (Gallon and Cullen, 2015).

**Figure 3.**
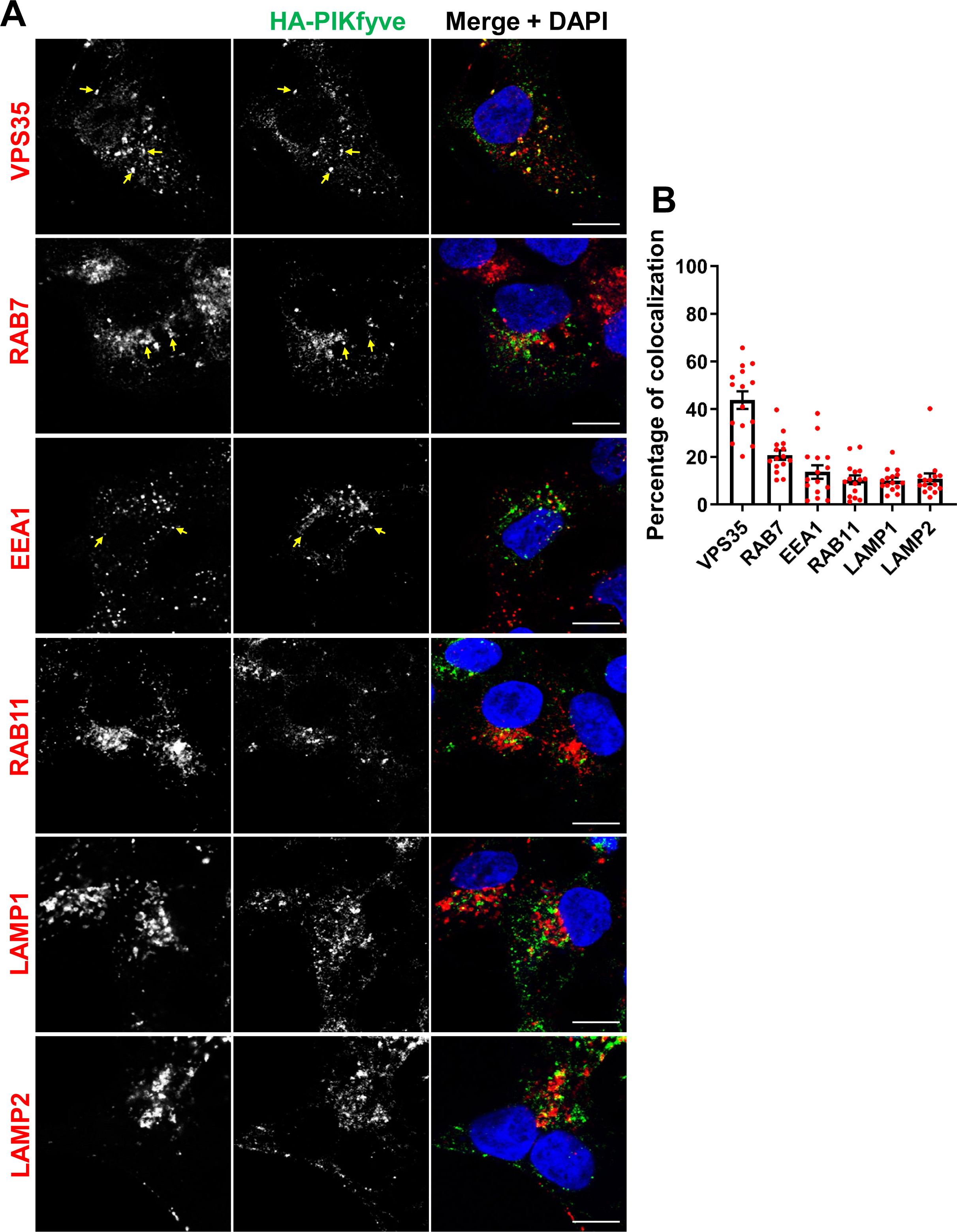
PIKfyve resides on early and late endosomes and exhibits the highest colocalization with VPS35 containing endosomes. A. HEK293 cells expressing 3xHA-endogenously tagged PIKfyve were fixed, permeabilized and immunostained with antibodies against the HA tag and with antibodies against proteins associated with the retromer (VPS35), early endosomes (EEA1), late endosomes (RAB7), recycling endosomes (RAB11) or lysosomes (LAMP1 and LAMP2). Arrows indicate examples of puncta showing colocalization. Bar-10 µm. B. The percentage of PIKfyve that colocalizes with the indicated proteins was determined using Mander’s colocalization coefficient from three independent experiments.

There was a modest colocalization of PIKfyve with RAB7 and EEA1, and some colocalization with RAB11, LAMP1 and LAMP2. The HA antibody uniquely recognized PIKfyve because no signal was observed in non-edited control HEK293 cells (Figure S3). Localization of PIKfyve to several endosomal compartments fits with our previous studies which found a significant pool of VAC14 on both early and late endosomes (Zhang et al., 2012) as well as studies showing that exogenous overexpressed PIKfyve partially co-localizes with early and late endosomes (McCartney et al., 2014b; Rutherford et al., 2006).

### Inhibition of PIKfyve delays the exit of internalized β1**-**integrin

β1-integrin cycles between the plasma membrane and endosomes (Moreno-Layseca et al., 2019). Since endogenous PIKfyve colocalizes with several endocytic compartments, we specifically tested the impact of PIKfyve inhibition on recycling of internalized β1-integrin back to the cell surface. To generate a pool of labeled, internalized β1-integrin, untreated HeLa cells were incubated for 1 h in the presence of an antibody against an extracellular epitope of β1-integrin. At the start of the recycling assay (0 time), cells were exposed to an acid wash to strip off any antibody bound to surface exposed β1-integrin. Cells were then treated with either apilimod or DMSO and further incubated for 15 or 30 min (Figure 4A-E). Notably, at 15 min, the amount of β1-integrin that recycled back to plasma membrane was significantly lowered by 0.58-fold in cells treated with apilimod as compared to DMSO-treated cells. Moreover, after 30 min of treatment, the apilimod-treated cells had 0.45-fold less surface β1-integrin compared with DMSO-treated cells (Figure 4K). In addition, we determined the internalized pool of β1-integrin that remained trapped in internal compartments. At the 15 and 30 min time points, we performed a second acid wash to remove antibodies attached to integrin that returned back to the cell surface (Figure 4F-J). Consistent with a defect in the return of β1-integrin back to the cell surface, inhibition of PIKfyve resulted in an increase in the internal, labeled, non-recycled pool of β1-integrin. In cells treated with apilimod for 15 min, the internal-non-recycled pool of β1-integrin was significantly higher by 20% compared with DMSO-treated cells, and this accumulation was 30% higher after 30 min of apilimod treatment (Figure 4L). These results indicate that PIKfyve is required for the recycling of β1-integrin from internal compartments to the plasma membrane.

**Figure 4:**
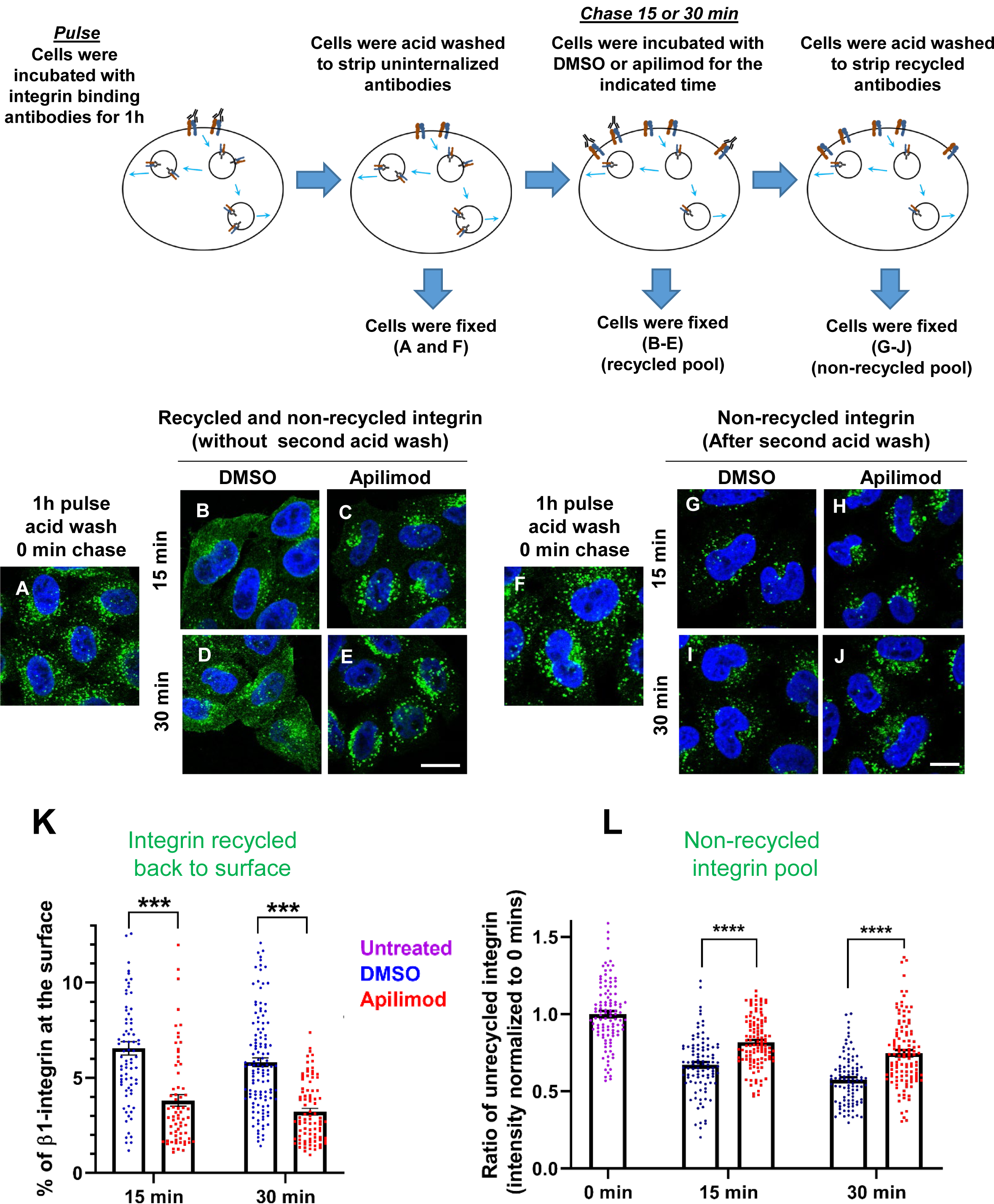
Inhibition of PIKfyve results in a defect in β1-integrin recycling. A-J. HeLa cells were incubated with β1-integrin antibody for 1 h at 37°C to allow the antibody-labeled integrin to internalize. Cells were then acid washed to remove surface β1-integrin bound antibodies, and were either fixed (A and F) or incubated with DMSO or 1 µM apilimod containing media at 37°C for the indicated times. Cells were either fixed (B-E)) or fixed after a second acid wash to remove antibodies that returned to the surface (G-J). Fixed cells were permeabilized and immunostained with Alexa-Fluor-488 conjugated anti-mouse secondary antibodies. Flow diagram (top) outlines the experiment. K. Surface levels of β1-integrin were inferred from the intensity of β1-integrin within 0.8 micron from the cell border. The levels of β1-integrin that recycled back to the surface (for images B-E) were quantified as percentage of the total labeled integrin. L. Intensity of non-recycled β1-integrin was quantified from cells treated as indicated in (G-J). All values were normalized to the average of the 0 min time point (F). Data presented as mean ± SE. Statistical significance from three independent experiments were analyzed using two-way ANOVA and Sidak’s multiple comparisons tests. (K-L). *** P<0.005 and **** P<0.001. Bar-10 µm.

### Inhibition of PIKfyve delays the exit of integrin from several endocytic compartments

β1-integrin traffics through several endocytic compartments. Following internalization into endosomes, a majority of β1-integrin is recycled back to the plasma membrane either by a slow (RAB11) or fast (RAB4) recycling pathway. To determine whether inhibition of PIKfyve delays the exit of β1-integrin from a specific type of endosome, we first generated a pool of labeled, internalized β1-integrin. HeLa cells were incubated for 1 h with antibodies that bind to surface-exposed β1-integrin to label the fraction that was internalized during this time frame. Then the remaining surface bound uninternalized antibodies were removed with a short acid wash (0 min, untreated). Consistent with previous studies, at 0 time, β1-integrin was predominantly present in EEA1-positive endosomes (untreated cells, Figure 5A-B). In addition, there was some colocalization of β1-integrin with RAB11, RAB4 and LAMP1-positive compartments.

**Figure 5:**
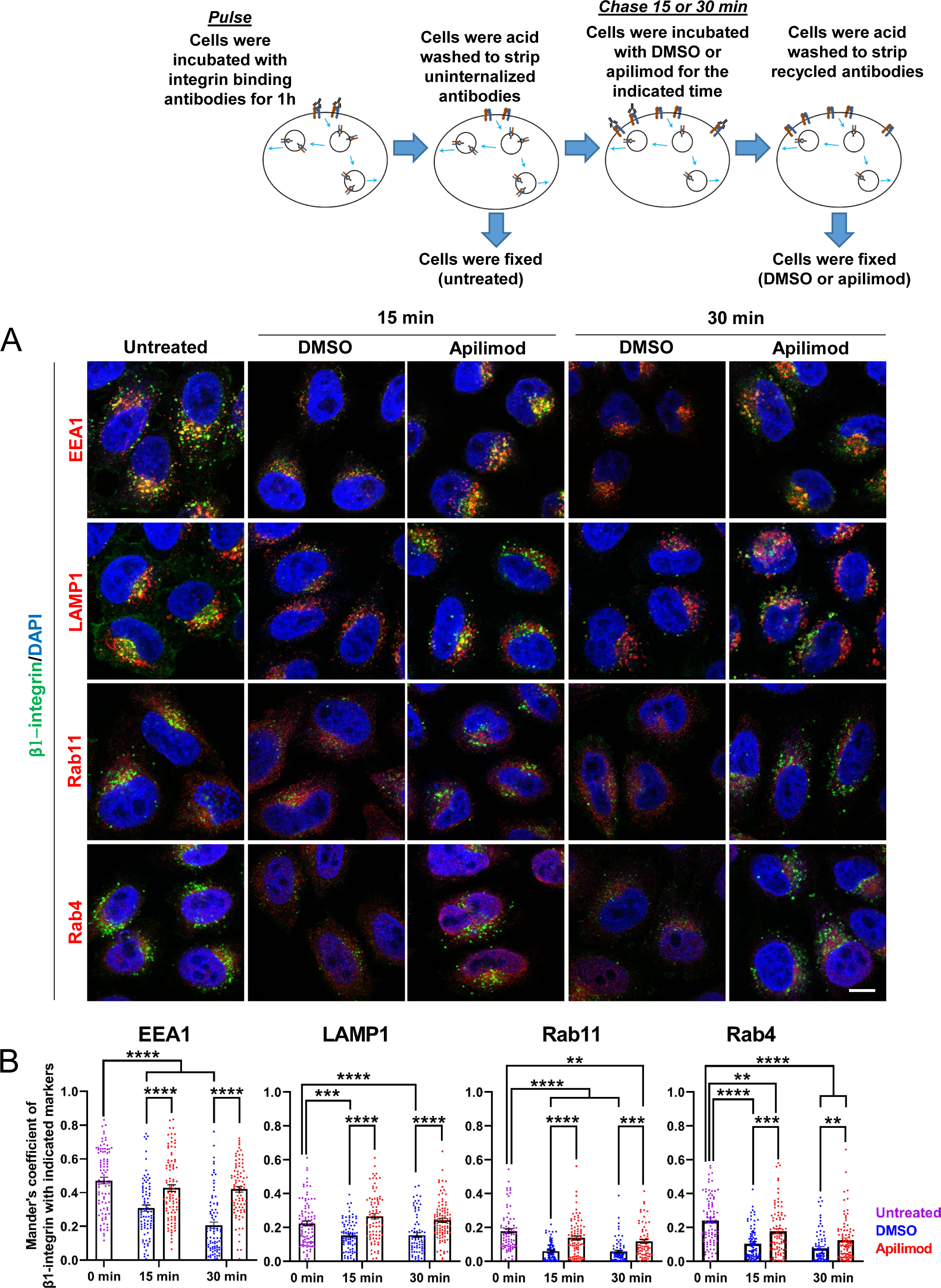
Inhibition of PIKfyve results in a defect in β1-integrin recycling from late endosomes, early endosomes and recycling endosomes. A. HeLa cells were incubated with β1-integrin antibody for 1 h at 37°C to allow antibody-labeled surface integrin to internalize and then acid washed to remove surface antibodies. Cells were either fixed (0 min, untreated) or treated with either DMSO or apilimod for the indicated time points. Cells were then acid washed, fixed and the localization of internalized β1-integrin was analyzed with well-established marker proteins: EEA1, LAMP1, RAB11 and RAB4. B. Colocalization of the internalized β1-integrin pool with endocytic markers was determined using Mander’s colocalization coefficient analysis. Data presented as mean ± SE. Statistical significance from three independent experiments was analyzed using two-way ANOVA and Sidak’s multiple comparisons tests. ** P<0.01, *** P<0.005 and **** P<0.001. Bar-10 µm.

To test whether PIKfyve inhibition altered the exit of internalized β1-integrin out of these compartments, cells were incubated with DMSO or apilimod for the indicated time points. Following treatment, an acid wash was performed to remove any antibody-bound integrin that was recycled back to the cell surface, and the remaining internalized pool was assessed. When compared with DMSO controls, PIKfyve inhibition resulted in significantly more labeled β1-integrin in each of the compartments that are part of its itinerary (Figure 5B). This suggests that PIKfyve activity is required for the exit of β1-integrin from several endocytic compartments.

Specifically, the amount of integrin present in EEA1 compartments in DMSO-treated cells decreased by 35% at 15 min and this amount further decreased to 56% at 30 min. In contrast, with PIKfyve inhibition, there was not a significant decrease in the amount of integrin present in EEA1 compartments at either 15 or 30 min. This indicates a delay in the exit of integrin from early endosomes. Similarly, in DMSO-treated cells, the amount of integrin present in LAMP1-positive compartments decreased approximately 32% at 15 and 30 min. However, following PIKfyve inhibition, there was no observable decrease in integrin in LAMP1 compartments. Together, these data show that PIKfyve inhibition causes a delay in the exit of β1-integrin from early and late endosomes. Note that, β1-integrin is recycled from both of these compartments (Moreno-Layseca et al., 2019).

Some of the β1-integrin in LAMP1 compartments could potentially be targeted for lysosomal degradation. However, at the time points measured, we did not see an impact of PIKfyve inhibition on the degradation of β1-integrin. The total level of β1-integrin was not significantly altered after apilimod treatment for 30 min (Figure S4).

We also tested whether PIKfyve activity is required for the trafficking of β1-integrin through either RAB4 or RAB11 compartments, which are part of the fast and slow recycling pathways, respectively. In cells treated with DMSO for 15 or 30 min, the amount of β1-integrin in RAB11-positive compartments decreased by approximately 66%. In contrast, during PIKfyve inhibition the amount of integrin in apilimod-treated cells had a more modest decrease of 22% and 34%, at 15 and 30 min of treatment respectively. The difference between DMSO and apilimod-treated cells suggested either a decrease in the rate of exit of integrin from RAB11 compartments or increased transport of integrin to RAB11 compartments from early endosomes. However, since the exit of integrin from early endosomes is also defective, the increase in integrin in RAB11 endosomes is likely due to defects in recycling of β1-integrin towards the plasma membrane.

There were similar defects in the trafficking of β1-integrin from RAB4 compartments. In cells treated with DMSO for 15 and 30 min, the amount of integrin remaining in RAB4 endosomes was significantly lower by 57% and 67% respectively. In comparison, following PIKfyve inhibition, the decrease in β1-integrin in RAB4 endosomes was only 27% at 15 min and 48% after 30 min. Thus, short-term inhibition of PIKfyve also slows the recycling of β1-integrin from RAB4 endosomes. Together, these studies suggest that PIKfyve plays a role in the recycling of β1-integrin from all endocytic compartments tested including early and late endosomes as well as fast and slow recycling endosomes.

### PIKfyve colocalizes with SNX17-Retriever-CCC-WASH complex proteins

β1-integrin recycling from endosomes is regulated by the sorting nexin, SNX17, the Retriever complex, the CCC complex, and the actin regulatory WASH complex (Chen et al., 2019; McNally and Cullen, 2018; Simonetti and Cullen, 2019; Wang et al., 2018). Loss of SNX17 inhibits the recycling of β1-integrin from endosomes (Bottcher et al., 2012; McNally et al., 2017; Steinberg et al., 2012).

β1-integrin is one of several cargoes that require SNX17, WASH, Retriever, and the CCC complex. To test whether PIKfyve is more generally required for SNX17, Retriever, CCC, and WASH complex-mediated trafficking from endosomes to the plasma membrane, we tested the impact of PIKfyve inhibition on two additional Retriever cargoes, α5-integrin and low density lipoprotein receptor-related protein 1 LRP1 (Farfan et al., 2013; McNally et al., 2017). We performed surface biotinylation assays, and found that inhibition of PIKfyve lowers β1-integrin, α5-integrin and LRP1 levels on the cell surface by approximately 50% each (Figure 6A-B). This finding suggests that PIKfyve regulates general SNX17-Retriever-CCC-WASH mediated recycling.

**Figure 6.**
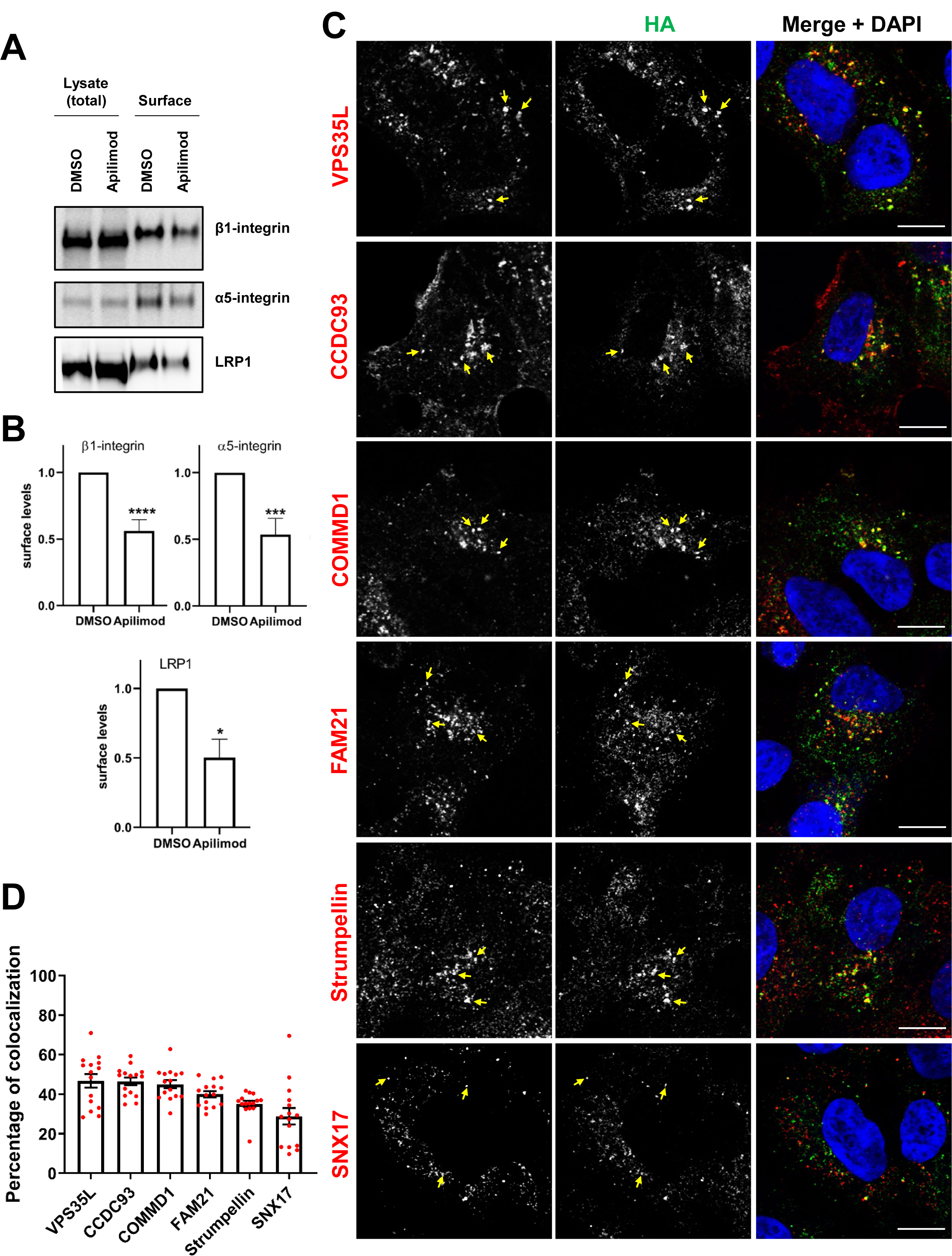
PIKfyve regulates the recycling of SNX17-Retriever-CCC-WASH cargoes and colocalizes with SNX17 and subunits of the Retriever, CCC and WASH complexes. A. HeLa cells were treated with DMSO or 1 µM apilimod for 1 h and then the levels of SNX17 cargoes, β1-integrin and α5-integrin were determined using a surface biotinylation assay. Surface biotinylation assay was similarly performed on HUH7 cells and LRP1 levels were measured. B. Quantification of western blots from three independent experiments of surface biotinylation experiments. C. HEK293 cells expressing 3xHA-endogenously tagged PIKfyve were fixed, permeabilized and incubated with antibodies against the HA tag and antibodies against either SNX17, the Retriever complex subunit (VPS35L), CCC complex subunits (CCDC93 and COMMD1) or WASH complex subunits (FAM21 and Strumpellin). Bar-10 µm. Arrows indicate examples of puncta showing colocalization. D. The percentage of PIKfyve colocalizing with the indicated proteins was determined using Mander’s colocalization coefficient analysis from three independent experiments. Data presented as mean ± SE. * P<0.05, *** P<0.005 and **** P<0.001.

We tested and found that PIKfyve colocalizes with SNX17 and the subunits of the WASH (Strumpellin and FAM21), Retriever (VPS35L) and CCC (COMMD1 and CCDC93) complexes (Figure 6C-D). Utilizing Mander’s coefficient, we quantified the fraction of the indicated proteins that overlap with endogenous PIKfyve-positive puncta, and observed a colocalization of 30-50% of endogenous PIKfyve with the proteins implicated in β1-integrin recycling. This colocalization was not observed in the non-edited control HEK293 cells (Figure S3). These data provide further support for the hypothesis that PIKfyve regulates β1-integrin recycling from endosomes via regulation of the SNX17-Retriever-CCC-WASH complex.

### PIKfyve regulates the localization of the CCC and Retriever complexes at endosomes

To gain mechanistic insight into how PIKfyve regulates β1-integrin recycling, we used HeLa cells and tested whether PIKfyve is required for the recruitment of SNX17 and/or Retriever-CCC-WASH complex subunits to endosomes. We tested colocalization of these proteins with VPS35-positive endosomes because both the retromer and Retriever pathways emerge from VPS35 containing endosomes (McNally et al., 2017; Singla et al., 2019). In addition, PIKfyve exhibits a strong colocalization with VPS35 (Figure 3). Strikingly, for each CCC and Retriever subunit tested, acute inhibition of PIKfyve for 30 min caused a significant decrease in their colocalization with VPS35 endosomes. The CCC subunits COMMD1 and COMMD5 were lowered by 22% and 20%, respectively (Figure 7, Figure S5 A-B). Importantly, the loss of the CCC proteins from VPS35 endosomes occurred over a relatively short time frame, 30 minutes. This may be due to some proteins in the CCC complex directly interacting with PI3,5P_2_ and/or PI5P. Importantly, COMMD1, COMMD7 and COMMD10 bind some phosphoinositides including PI3,5P_2_ and in some cases PI5P in in vitro assays (Healy et al., 2018).

**Figure 7.**
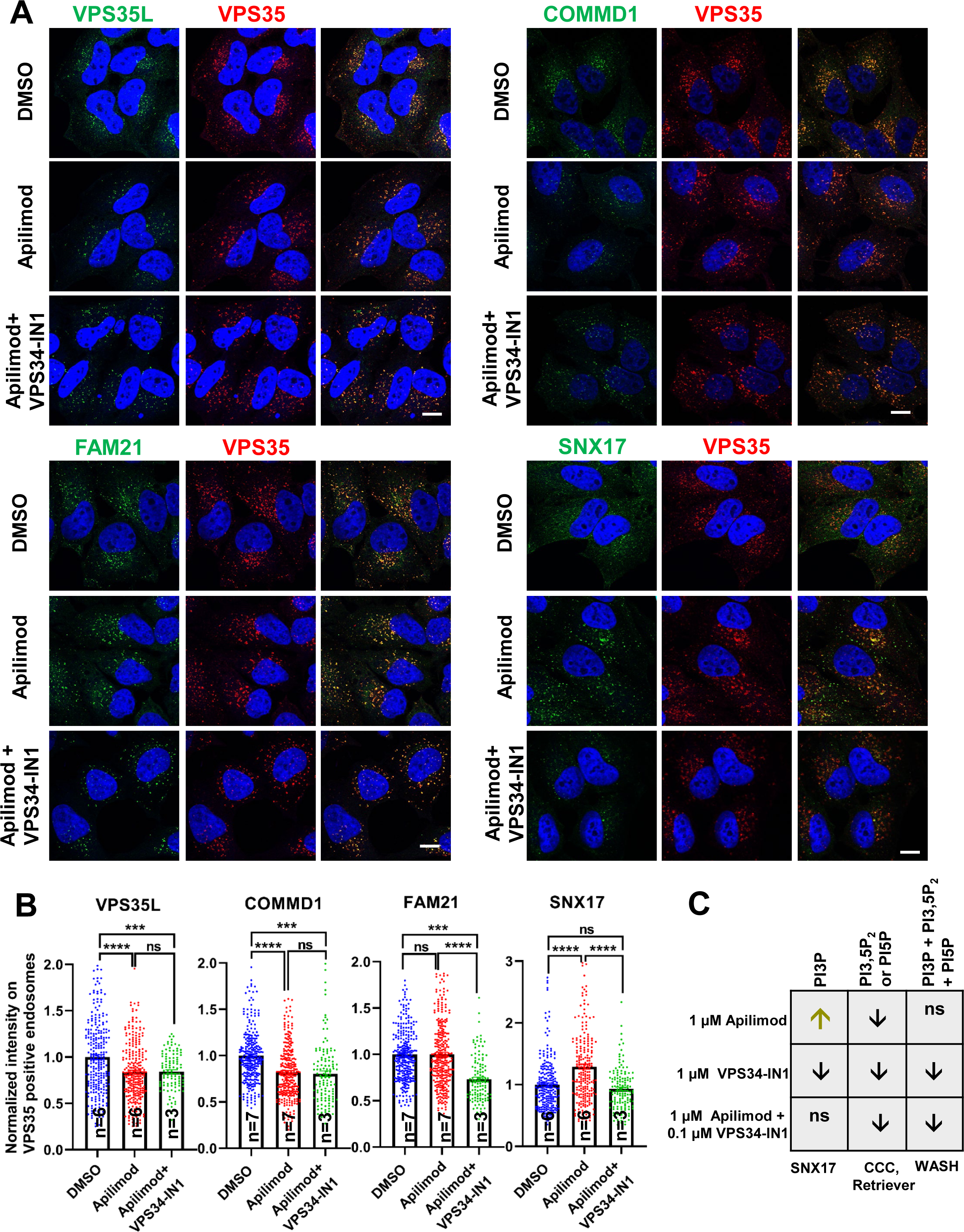
CCC and Retriever complexes require PI3,5P_2_ and/or PI5P to bind to endosomes. A. HeLa cells treated with either DMSO or 1 µM apilimod or co-treated with 1 µM apilimod and 0.01 µM VPS34-IN1 for 30 min were fixed, permeabilized and co-stained with antibodies against VPS35 (A-D) and antibodies against either VPS35L, COMMD1, FAM21 or SNX17. B. The intensity of VPS35L, COMMD1, FAM21 and SNX17 on VPS35-positive endosomes was quantified and values were normalized to the corresponding average intensity of the DMSO treatment cohort. C. Summary of the effect of inhibitor treatment on phosphoinositide lipid levels and the endosomal localization of SNX17, WASH, CCC and Retriever subunits. Khaki and black arrows indicate an increase or decrease, respectively. Data presented as mean ± SE. Statistical significance from three or more independent experiments as indicated within bar graph were analyzed using one-way ANOVA and Tukey post hoc tests. *** P<0.005 and **** P<0.001, and ns, not significant. Bar-10 µm.

The Retriever subunit, VPS35L was lowered by 17% (Figure 7A-B). We were unable to test VPS26C, the other subunit unique to Retriever, because we have not identified an antibody suitable for immunofluorescence. The reliance of the Retriever complex on PIKfyve may either be due to direct binding to PI3,5P_2_ and/or PI5P, or may be due to a requirement for the presence of the CCC complex on endosomes.

In contrast with the partial loss of CCC and Retriever proteins from membranes, acute inhibition of PIKfyve resulted in a 27% increase in SNX17 on VPS35 positive endosomes (Figure 7A-B). The increased recruitment of SNX17 may be due to an elevation in PI3P that occurs during inhibition of PIKfyve (Zolov et al., 2012). The WASH complex subunit, FAM21 remained unchanged.

To further determine whether the effects observed during PIKfyve inhibition were due to elevation in PI3P or lowering PI3,5P_2_ or PI5P, we sought to determine the enzyme that generates the PI3P pool that recruits SNX17. In mouse embryonic fibroblasts, approximately, two-thirds of the PI3P pool comes from VPS34 (Devereaux et al., 2013; Ikonomov et al., 2015). Note that inhibition of VPS34 also lowers PI3,5P_2_ and PI5P (Figure S6) and see also (Devereaux et al., 2013; Ikonomov et al., 2015). This is because PI3,5P_2_ is synthesized from PI3P (Sbrissa et al., 1999), and PI5P can be synthesized from PI3,5P_2_ (Zolov et al., 2012).

To further test a role for PI3P in the recruitment of SNX17, we inhibited VPS34 with VPS34-IN1, and found that there was significantly less SNX17 on VPS35 endosomes. In addition, we observed less VPS35L, COMMD1 (Figure S7), and COMMD5 (Figure S5). These changes are likely due to lower levels of PI3,5P_2_ and PI5P, as inhibition of PIKfyve with apilimod also lowered the levels of these proteins on VPS35 endosomes. The levels of FAM21 were not impacted and remained similar to untreated cells.

To determine effects due to lowering PI3,5P_2_ and PI5P, under conditions where PI3P levels are not elevated, we tested and found that a combination of 1 μM apilimod, and 0.1 µM VPS34-IN1, resulted in lower PI3,5P_2_ and PI5P, but normal cellular levels of PI3P (Figure S8). Note that the methods used here to measure phosphoinositides, report on total cellular levels and would not detect a change at a specific membrane domain.

We tested and found that lowering PI3,5P_2_ and PI5P under conditions without detectable changes in PI3P, resulted in no change in SNX17 recruitment. However, there was less VPS35L, COMMD1, and FAM21 on VPS35 endosomes. A summary of trends in phosphoinositide levels with each treatment are shown in Figure 7C. These studies indicate that SNX17 is recruited by PI3P, that VPS35L (Retriever) and COMMD1 (CCC) are either directly or indirectly recruited by PI3,5P_2_ and/or PI5P, and that FAM21 can be recruited by PI3P and/or PI3,5P_2_ and PI5P. Consistent with this finding, FAM21 binds multiple phosphorylated phosphoinositide lipids in vitro (Jia et al., 2010). Together, these findings suggest that that the SNX17-Retriever-CCC-WASH recycling pathway may be ordered by phosphoinositide conversion, where PI3P recruits SNX17 to WASH complex-containing endosomes, and PI3P-dependent recruitment of PIKfyve generates PI3,5P_2_ and/or PI5P which recruits the CCC and Retriever complexes.

To further probe how PIKfyve recruits the CCC complex, we focused on COMMD1, a CCC subunit which functions as an obligate dimer and binds multiple phosphorylated phosphoinositide lipids including PI3,5P_2_ and PI5P in vitro (Healy et al., 2018). Mutation of residues that comprise a basic patch on COMMD1, R133Q, H134A and K167A (COMMD1-QAA), abolished the ability of COMMD1 to bind phosphoinositide lipids in vitro (Healy et al., 2018). However, this mutant bound membranes in cells, although there were no further tests of function. Note that in this mutant, two of the basic residues were substituted with alanine, which is hydrophobic and could potentially cause non-specific sticking to cellular membranes. Thus, we mutated the same sites as follows R133E, H134Q and K167E (COMMD1-EQE). Moreover, we noted that these residues are on one side of a basic pocket, with K129 nearby, and R120 from the other monomer nearby on the other side of the pocket (Figure 8A). Consequently, we also generated COMMD1-R120E, K129D, R133E, H134Q, K167E, (COMMD1-EDEQE). Importantly all the mutated residues are predicted to be surface exposed, and these changes should not change the overall structure of COMMD1. We expressed these mutants as well as the original mutant in COMMD1-/- knock-out cells and found that as previously reported, there was no statistically significant difference in the binding of COMMD1 and the original COMMD1-QAA mutant to endosomes. In contrast, both the COMMD1-EQE and COMMD1-EDEQE mutants exhibited defects in their association with VPS35 endosomes, which are statistically significant, 0.14 and 0.22-fold, respectively (Figure 8B-C). These findings indicate that the phosphoinositide binding site of COMMD1 plays a role in its association with membranes, and strongly suggests that phosphoinositide binding is important for COMMD1 function.

**Figure 8.**
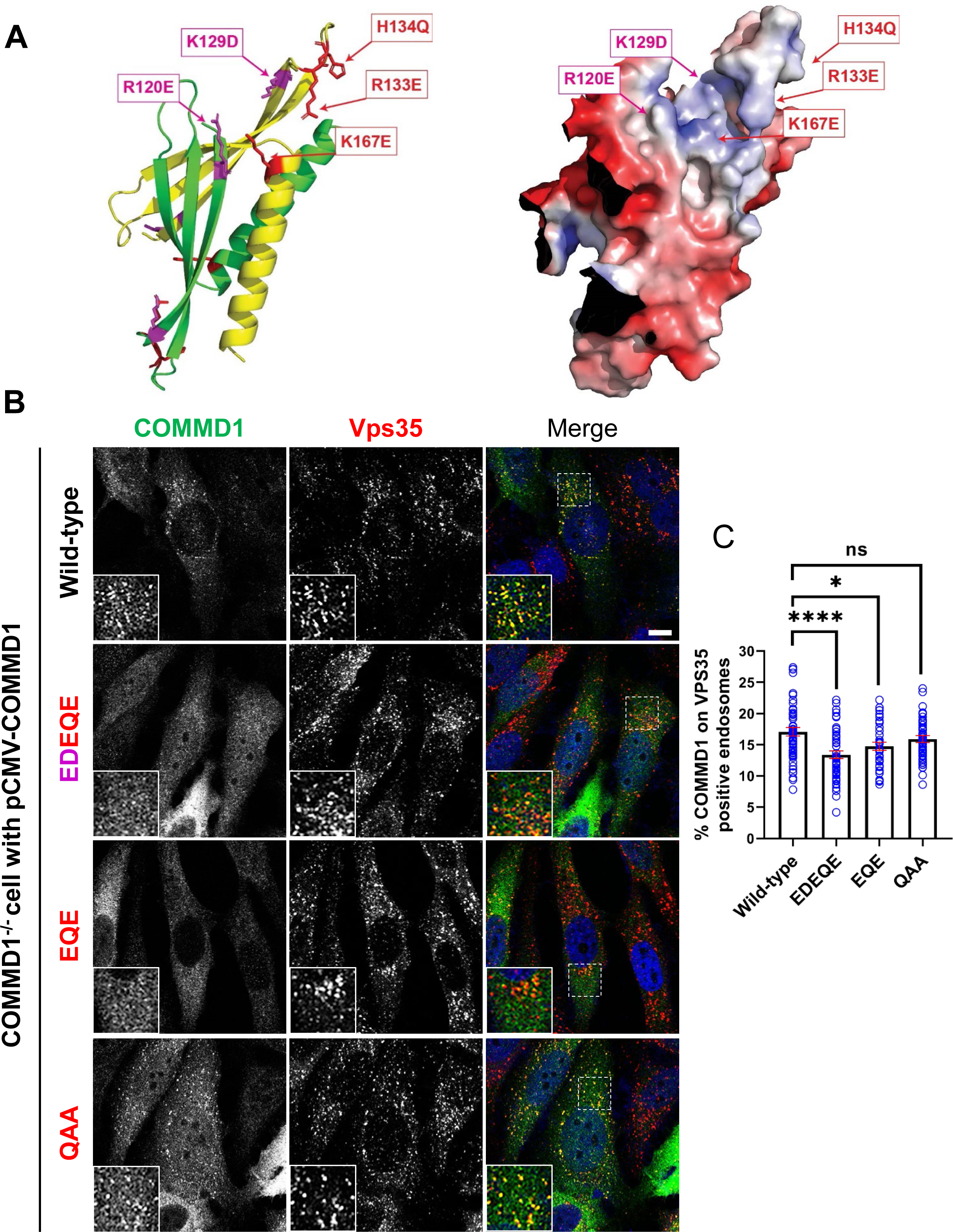
Mutation of the putative phosphoinositide binding site impairs COMMD1 localization to VPS35-positive endosomes. A. Ribbon and space filling models of the COMMD domain of COMMD1 modeled on COMMD9 (PDB: 6BP6). Positively charged residues within the predicted PPI binding site are indicated. B-C. COMMD1 knock-out HeLa cells were transiently transfected with either wild type COMMD1 or COMMD1 mutants (EDEQE, EQE and QAA), then fixed, permeabilized and co-stained with antibodies against COMMD1 and VPS35. EDEQE: R120E/ K129D/ R133E/ H134Q/ K167E. EQE: R133E/ H134Q/ K167E, and QAA: R133Q/ H134A/ K167A. The percent of the total COMMD1 residing on VPS35-positive endosomes was quantified. Data presented as mean ± SE. Statistical significance from three independent experiments were analyzed using one-way ANOVA and Dunnett’s post hoc test. * P<0.05, and **** P<0.001, and ns not significant. Bar-10 µm.

To determine the functional significance of COMMD1 binding to phosphoinositide lipids in β1-integrin recycling, we expressed wild-type and the COMMD1-EDEQE mutant in COMMD1 knock-out cells. Cells were incubated with antibodies against β1-integrin for 1 h to allow the antibodies to internalize, then the remaining surface bound antibodies were removed with an acid wash. Immunofluorescence localization of the internalized pool of β1-integrin revealed no significant difference in β1-integrin internalization in untransfected cells or cells transfected with wild-type COMMD1 or the COMMD1-EDEQE mutant (Figure 9A, 9C). Cells with internalized β1-integrin were then incubated in serum containing media for 1 hour and the non-recycled pool of β1-integrin was assessed following a second acid wash (Figure 9B, 9D). Cells expressing wild-type COMMD1 had 27% less non-recycled β1-integrin compared to untransfected cells, which indicates the maximum restoration of β1-integrin that would occur with wild-type COMMD1. Importantly, the COMMD1-EDEQE mutant failed to correct the recycling defect. The difference between cells transfected with the COMMD1-EDEQE mutant vs. non-transfected cells, was 9%, which is not significantly different. Note that the difference between the COMMD1-EDEQE mutant and wild-type COMMD1 was significantly different. Together these results demonstrate that COMMD1 binding to phosphoinositide lipids is required for β1-integrin recycling.

**Figure 9.**
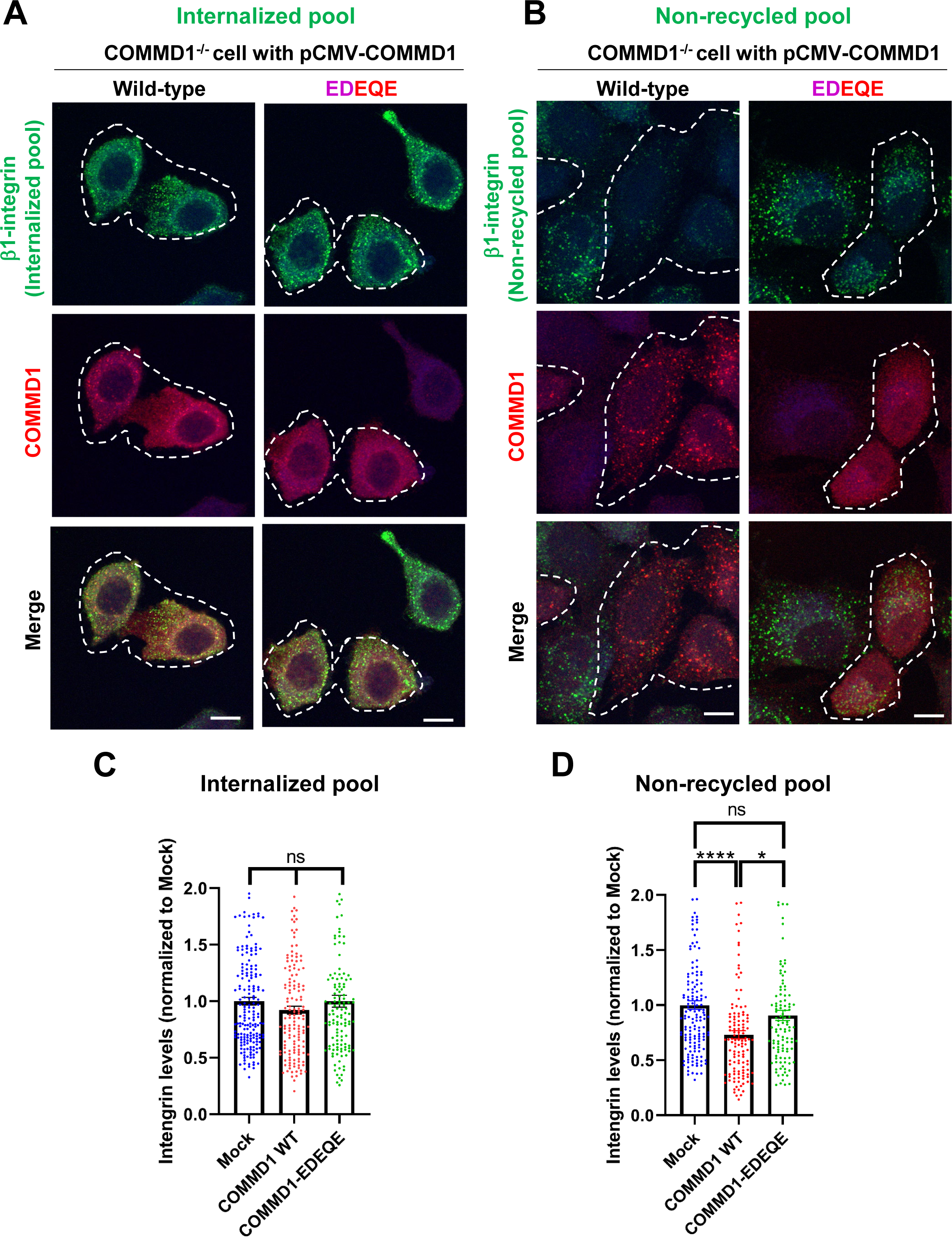
Mutation of the putative phosphoinositide binding site on COMMD1 delays recycling of β1-integrin. A-B COMMD1 knock-out HeLa cells were either untransfected (mock) or transiently transfected with wild type COMMD1 or the COMMD1-EDEQE (R120E/ K129D/ R133E/ H134Q/K167E) mutant for 16 hours. Cells were then incubated with anti β1-integrin antibodies for 1 h in serum containing media. Cells were acid washed. Cells were either fixed (A and C) or incubated with serum containing media for 1 hour. Cells were acid washed once again and fixed (B and D). C-D The intensity of integrin was quantified from four (D) or five (C) independent experiments and values were normalized to the corresponding average intensity of the mock treatment cohort. Data presented as mean ± SE. Statistical significance were analyzed using one-way ANOVA and Dunnett’s (C) or Turkey’s (E) post hoc test. * P<0.05, *** P<0.005 and ns not significant. Bar-10 µm. Transfected cells highlighted with white dotted lines.

In addition to regulation of the SNX17-Retriever-CCC-WASH recycling pathway, PIKfyve could potentially play a role in β1-integrin recycling via control of endosome associated actin. A previous study revealed that PIKfyve negatively regulates cortactin, which in turn causes excessive Arp2/3 activity and hyperaccumulation of actin on endosomes (Hong et al., 2015). In those studies, PIKfyve was inhibited for 2 hours. However, in the shorter time-frame of PIKfyve inhibition used in this study, we did not find any changes in the amount of cortactin that localizes to endosomes (Figure S10).

## Discussion

The best characterized roles of PIKfyve are its functions on lysosomes. Inhibition or depletion of PIKfyve results in enlarged lysosomes and late endosomes (de Araujo et al., 2020; Dove et al., 2009; Ho et al., 2012; McCartney et al., 2014a; Shisheva, 2012), and indeed, PIKfyve regulates multiple pathways that contribute to lysosome function. These include the regulation of multiple lysosomal ion channels (Chen et al., 2017; Dong et al., 2010; Fine et al., 2018; She et al., 2018; She et al., 2019; Wang et al., 2017), as well as roles in the fission and reformation of lysosomes and related organelles (Bissig et al., 2017; Choy et al., 2018; Krishna et al., 2016; Yordanov et al., 2019).

While the most apparent defects in cells following loss of PIKfyve activity are enlarged lysosomes, the multiple pleiotropic defects observed strongly suggest that PIKfyve plays key roles elsewhere in the cell. As an approach to gain mechanistic insight into roles for PIKfyve that are not lysosome-based, we sought mechanistic insight into roles for PIKfyve in cell migration.

The studies reported here reveal that PIKfyve has roles at endosomes, and has a direct role in the regulation of the SNX17-Retriever-CCC-WASH complex. Moreover, endogenously-tagged PIKfyve extensively colocalizes with SNX17, Retriever, CCC and WASH complexes. PIKfyve has a robust 44% colocalization with VPS35, a subunit of the retromer complex, as well as with the Retriever subunit, VPS35L. In addition, it is likely that PIKfyve specifically localizes to endosomes that are actively engaged in membrane transport. Early endosomes undergo rapid conversion to late endosomes (Rink et al., 2005), and in this process is linked to retromer and Retriever-based transport which occur from the same membrane subdomains (McNally et al., 2017; Singla et al., 2019). Moreover, RAB5 and RAB7 act in concert to regulate retromer recruitment to endosomes (Seaman et al., 2009). Thus, the hypothesis that PIKfyve is specifically localized to endosomal compartments involved in recycling fits with the observation that the best colocalization of PIKfyve is with VPS35, followed by good colocalization with EEA1 and RAB7.

Our mechanistic analysis of roles of PIKfyve with the SNX17-Retriever-CCC-WASH complexes, relied primarily on acute inhibition, which is likely to reveal pathways where PIKfyve plays a direct role. In addition, these studies were aided by utilization of a hyperactive allele of PIKfyve. Importantly, activation of PIKfyve had the opposite effect of PIKfyve inhibition, which provides additional evidence that PI3,5P_2_ and/or PI5P play direct roles in the SNX17-Retriever-CCC-WASH pathway, and in β1-integrin recycling.

Results reported here, together with earlier studies reveal that the SNX17-Retriever-CCC-WASH recycling pathway is ordered by changes in phosphoinositide lipids as well as a web of protein-protein interactions (Figure 10). VPS34, which resides on endosomes (Christoforidis et al., 1999), provides the PI3P that recruits SNX17 (Figure 7, Figure S7). The generation of PI3P recruits SNX17 (Chandra et al., 2019; Jia et al., 2014) and the WASH complex (Jia et al., 2010), and SNX17 binds its cargoes (Bottcher et al., 2012; Steinberg et al., 2012). The generation of PI3P also recruits PIKfyve, via its FYVE domain (Stenmark et al., 2002), which initiates the production of PI3,5P_2_. The generation of PI3,5P_2_ and/or PI5P then recruits the CCC complex (Figure 7). The CCC complex also binds the WASH complex via direct interaction of CCDC93 with the WASH subunit, FAM21 (Phillips-Krawczak et al., 2015). The Retriever complex may also be recruited directly by PI3,5P_2_ and/or PI5P (Figure 7), and/or may indirectly require PIKfyve activity to recruit the CCC complex, since the Retriever complex interacts with and requires the CCC complex to bind to endosomes (McNally et al., 2017; Phillips-Krawczak et al., 2015; Singla et al., 2019). In addition, the Retriever subunit, VPS26C interacts directly with SNX17 (Farfan et al., 2013; McNally et al., 2017).

**Figure 10.**
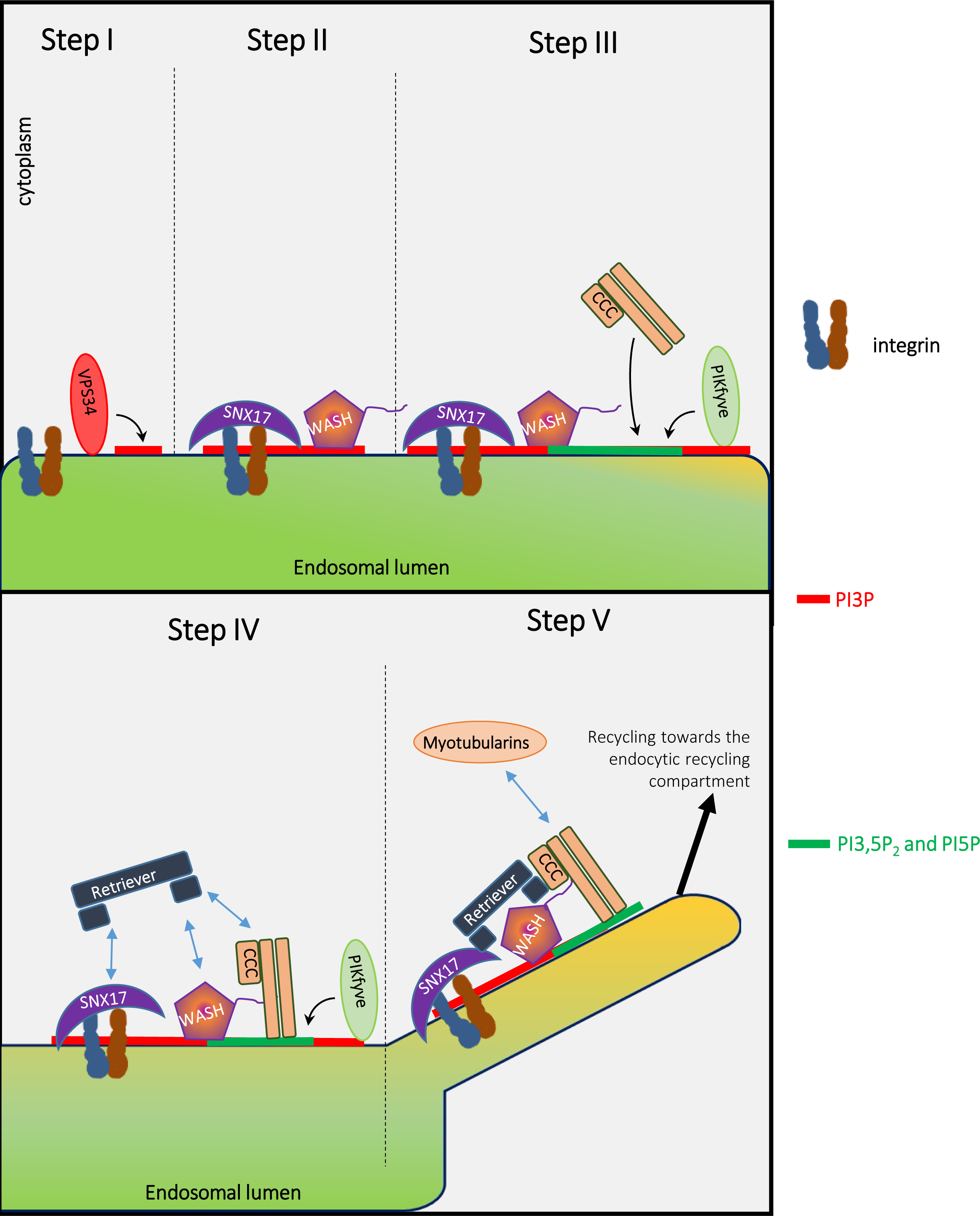
Model: VPS34 and PIKfyve regulate recycling of integrins from endosomes to the plasma membrane via promoting the ordered assembly of SNX17, WASH, Retriever and CCC complexes. The PI3P on cargo-containing endosomes is generated by Vps34 (Figures 7, S7). PI3P recruits SNX17 (Chandra et al., 2019; Jia et al., 2014), and SNX17 binds its cargo (Bottcher et al., 2012; Steinberg et al., 2012). The generation of PI3P likely recruits PIKfyve, via its FYVE domain (Stenmark et al., 2002), which initiates the production of PI3,5P_2_ and PI5P. The presence of the WASH complex relies on the generation of either of PI3P, PI3,5P_2_ and/or PI5P. The generation of PI3,5P_2_ and/or PI5P by PIKfyve then recruits the CCC complex (Figure 7). The CCC complex also binds the WASH complex via direct interaction of CCDC93 with the WASH subunit, FAM21 (Phillips-Krawczak et al., 2015). The Retriever complex may also be recruited directly by PI3,5P_2_ or PI5P, and/or may indirectly require PIKfyve activity to recruit the CCC complex. The Retriever complex interacts with and requires the CCC complex to bind to endosomes (McNally et al., 2017; Phillips-Krawczak et al., 2015; Singla et al., 2019). In addition, the Retriever subunit VPS26C interacts directly with SNX17 (Farfan et al., 2013; McNally et al., 2017). Importantly, while SNX17 and WASH are necessary, they are not sufficient for the recruitment of either Retriever or CCC (Figure 7, apilimod treatment). Recruitment of the Retriever and CCC complexes requires PIKfyve. Furthermore, the CCC subunit CCDC22, in turn recruits MTMR2, which is required for late steps in this recycling pathway (Singla et al., 2019). Importantly, recruitment of MTMR2 lowers PI3P and PI3,5P_2_ (Singla et al., 2019). Together, these studies suggest PI3P and PI3,5P_2_ coordinate the SNX17-Retriever-CCC-WASH pathway. PI3P in required for initiation, PI3,5P_2_ and/or PI5P act in the middle steps, and MTMr2, which removes PI3P and PI35P_2_, acts late in the pathway. Once the SNX17-Retriever-CCC-WASH complex assembles with cargo, the WASH complex mediates actin nucleation (Derivery et al., 2009; Gomez and Billadeau, 2009) and SNX17 recruits EHD1 (Dhawan et al., 2020) to enable fission of cargo containing membranes for efficient recycling.

The studies reported here also show that SNX17 and the WASH complex can bind to endosomes without the Retriever and CCC complexes. However, endosomes that contain SNX17 and the WASH complex are not sufficient for the recruitment of either the Retriever or CCC complexes. PIKfyve activity is also needed for recruitment of Retriever and CCC complexes. Furthermore, the CCC subunit CCDC22, recruits MTMR2, which is required for late steps in this recycling pathway (Singla et al., 2019). Importantly, we found that recruitment of MTMR2 lowers both PI3P and PI3,5P_2_ (Singla et al., 2019). Thus, the SNX17-Retriever-CCC-WASH pathway may be ordered in part via an initiation step that involves PI3P, middle steps that require PI3,5P_2_ and/or PI5P and a late step via MTMR2 that removes PI3P and PI3,5P_2_. Finally, once the SNX17-Retriever-CCC-WASH complex assembles with cargo, the WASH complex mediates actin nucleation (Derivery et al., 2009; Gomez and Billadeau, 2009) and SNX17 recruits EHD1 (Dhawan et al., 2020) to enable fission of cargo containing membrane for efficient recycling.

This new role for PIKfyve in endocytic recycling may partially explain a recent study which showed that treatment of cells with 100 nM apilimod for 16 h resulted in a reduction in steady-state levels of exogenously expressed, FLAG-TGFβ-R2 at the cell surface. Note that the recycling pathway utilized by TGFβ-R2 is currently unknown (Cinato et al., 2021).

Previous studies have also revealed roles for PIKfyve on endosomes. The PIKfyve pathway plays a role in the formation of Stage I melanosomes, which are derived from early endosomes (Bissig et al., 2019). Furthermore, we and others previously showed that knock-down of the PIKfyve pathway causes a defect in recycling of CI-MPR from endosomes to the trans-Golgi network (de Lartigue et al., 2009; Rutherford et al., 2006; Zhang et al., 2007), although the mechanistic basis of this defect remains unknown. Moreover, a recent study revealed that SNX11, which plays a role in the delivery of some cargoes from endosomes to lysosomes, binds to PI3,5P_2_ (Xu et al., 2020). In addition, PIKfyve inhibition caused the accumulation of the tight junction proteins, claudin1 and claudin2 into endosomes and delayed the formation of epithelial permeability barrier (Dukes et al., 2012). Interestingly, PIKfyve inhibition did not affect the surface localization of claudin4. This suggests that PIKfyve regulates specific recycling pathways. Additionally, heterologous studies in *Xenopus laevis* oocytes, suggested a connection between PIKfyve activity and RAB11 endosomes that regulate endocytic recycling (Seebohm et al., 2012; Seebohm et al., 2007). Furthermore, we previously found that PIKfyve provides acute regulation of the levels of the α-amino-3-hydroxy-5-methyl-4-isoxazolepropionic acid receptor (AMPAR) at neuronal postsynaptic sites (McCartney et al., 2014b; Zhang et al., 2012). Those studies suggested that PIKfyve is a negative regulator of AMPAR recycling, which is the opposite of PIKfyve providing positive regulation of α5-integrin, β1-integrin and LRP1 recycling. The difference may be that while these latter proteins traffic via the SNX17-Retriever-CCC-WASH pathway, AMPAR recycles via the SNX27-retromer pathway (Hussain et al., 2014). Thus, PIKfyve may differentially regulate these two pathways. Alternatively, the role of PIKfyve in recycling from endosomes to the plasma membrane in neurons may be different from the role of PIKfyve in other cell-types.

In addition to regulating surface levels of β1-integrin, PIKfyve likely regulates cell migration via control of additional pathways. Previous studies suggested that PIKfyve may also regulate cell migration via the activation of the Rac1-GTPase (Dayam et al., 2017; Oppelt et al., 2014). PIKfyve regulation of Rac1 activation and integrin recycling may occur in parallel. In addition, there is extensive cross-talk between Rho GTPases and focal adhesions (Vitali et al., 2019), thus integrin trafficking and Rac1 activation may be linked with each other.

Apilimod, the PIKfyve inhibitor used in these studies, has recently been proposed as a drug to investigate further for treatment COVID-19 (Kang et al., 2020; Ou et al., 2020; Riva et al., 2020), and also blocks the infection of Ebola virus in cells (Nelson et al., 2017; Qiu et al., 2018). Apilimod may block SARS-CoV-2 entry via loss of positive regulation of TPC2, a downstream target of PIKfyve that resides on lysosomes (Ou et al., 2020). The new studies presented here suggest an additional potential mechanism and indicate that the effect of apilimod on surface levels of angiotensin-converting enzyme 2 (ACE2) and Neuropilin-1 should also be investigated. ACE2 and Neuropilin-1, levels at the plasma membrane are regulated. Importantly, each have been proposed to serve as receptors for SARS-CoV-2 (Daly et al., 2020; Wrapp et al., 2020). Moreover, a genome wide screen to identify proteins that regulate entry of SARS-CoV2 identified subunits of the CCC, Retriever and WASH complexes (Zhu et al., 2021). Together, the potential of apilimod as an antiviral drug heightens the urgency of determining the multiple cellular functions of PIKfyve, and the elucidation of pathways that are impacted by acute inhibition of PIKfyve.

## Acknowledgements

We thank Bagyasree Jambunathan and Madison Tluczek for their assistance in image analysis. We thank Aaron D. Cohen, as well as members of the Weisman lab for helpful discussions. We thank Loren Brown and Alan Smrcka (University of Michigan-Ann Arbor) for generously providing primary neonatal cardiac fibroblasts. We Thank David Ginsburg Lab (University of Michigan-Ann Arbor) for providing HUH7 cells. This work was supported by R01-NS099340 and R01 NS064015 to LSW, R01-DK107733 to EB and DDB, and by the University of Michigan Protein Folding Diseases Fast Forward Initiative. SSPG was supported in part by a postdoctoral fellowship from the American Heart Association, 14POST20480137. GL was supported in part by a postdoctoral fellowship from the American Heart Association, 19POST34450253.

## Author contributions

SSPG, GL, PRR and LSW conceived and designed the project. DDB, EB and HT provided unpublished data that aided in the development of the project. MAS was involved in the conception and design of the microscopy experiments. NS proposed putative phosphoinositide binding residues on COMMD1 and generated the corresponding figure. SSPG, GL and PRR performed experiments, SSPG, GL, PRR and LSW analyzed the data. DDB provided cells and antibodies. SSPG, GL, PRR and LSW wrote the manuscript, and all authors read, revised, and approved the manuscript.

## Materials and Methods

### Reagents

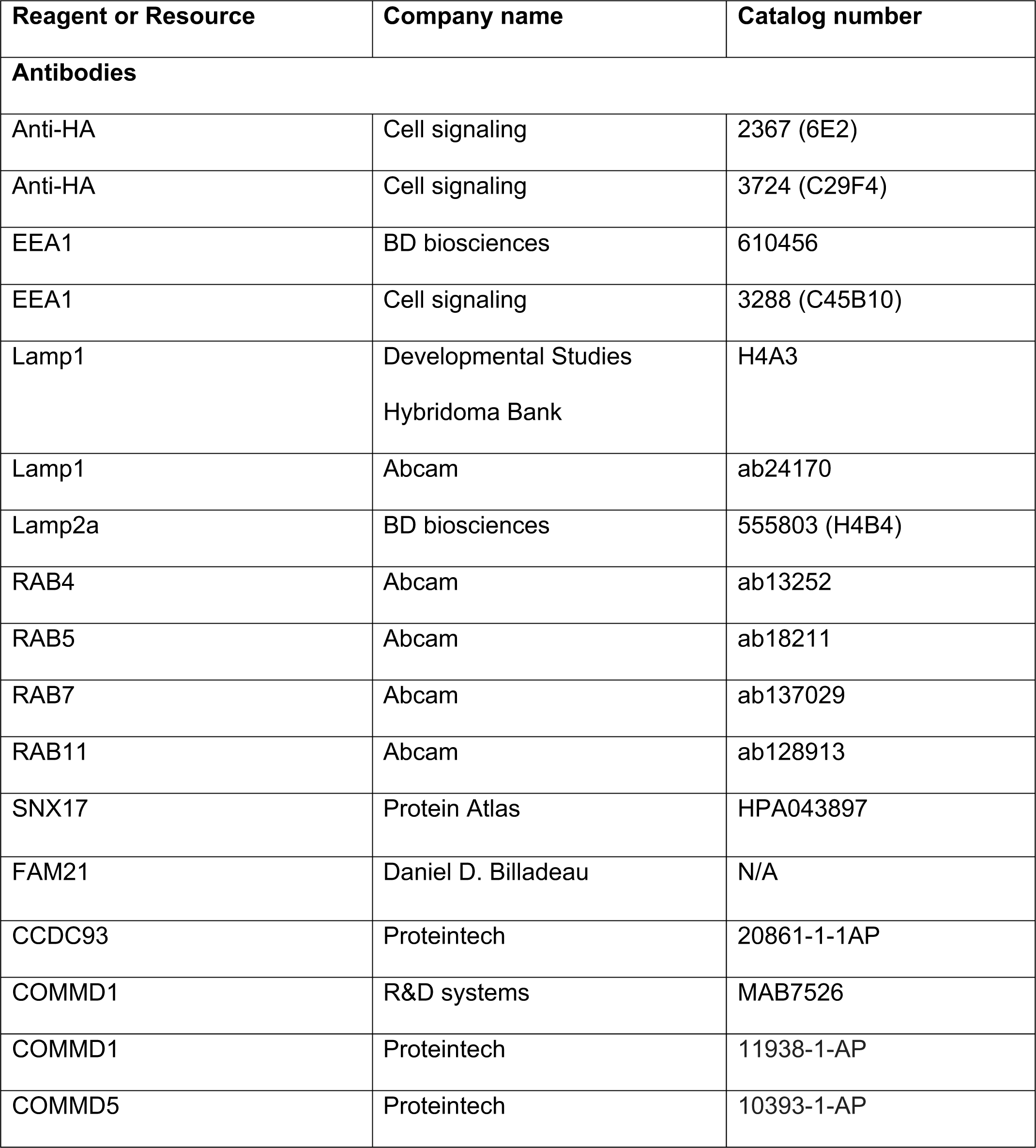
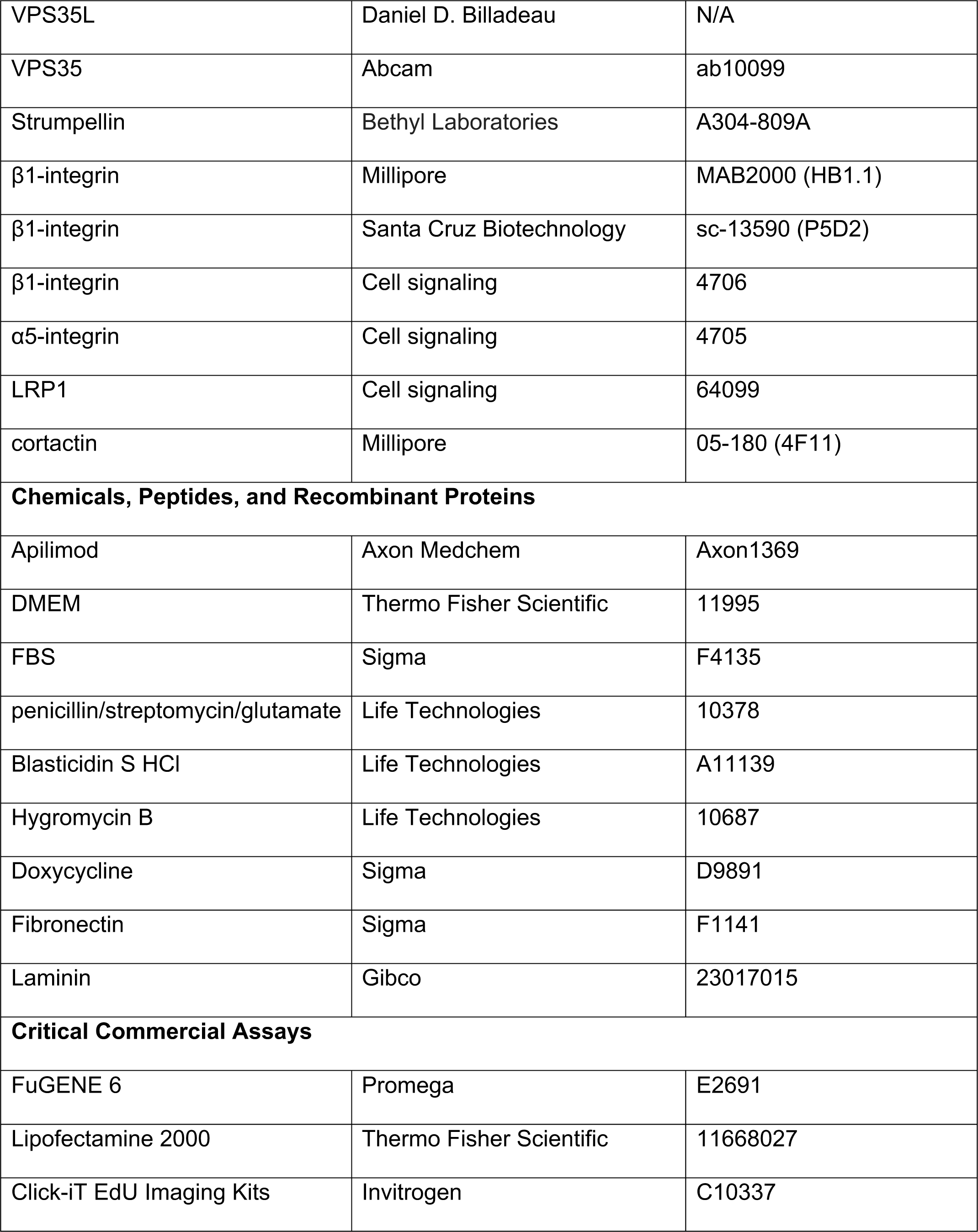
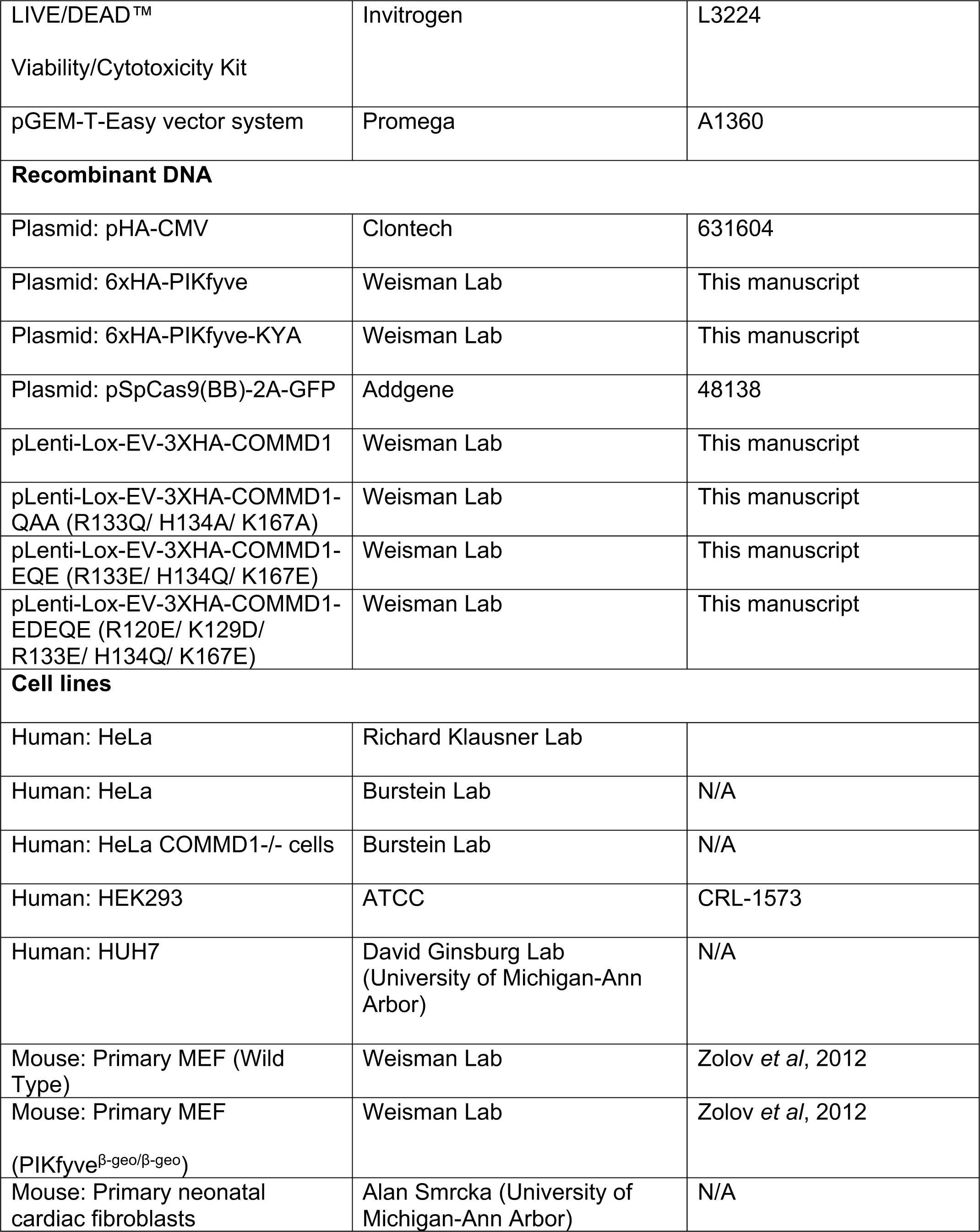
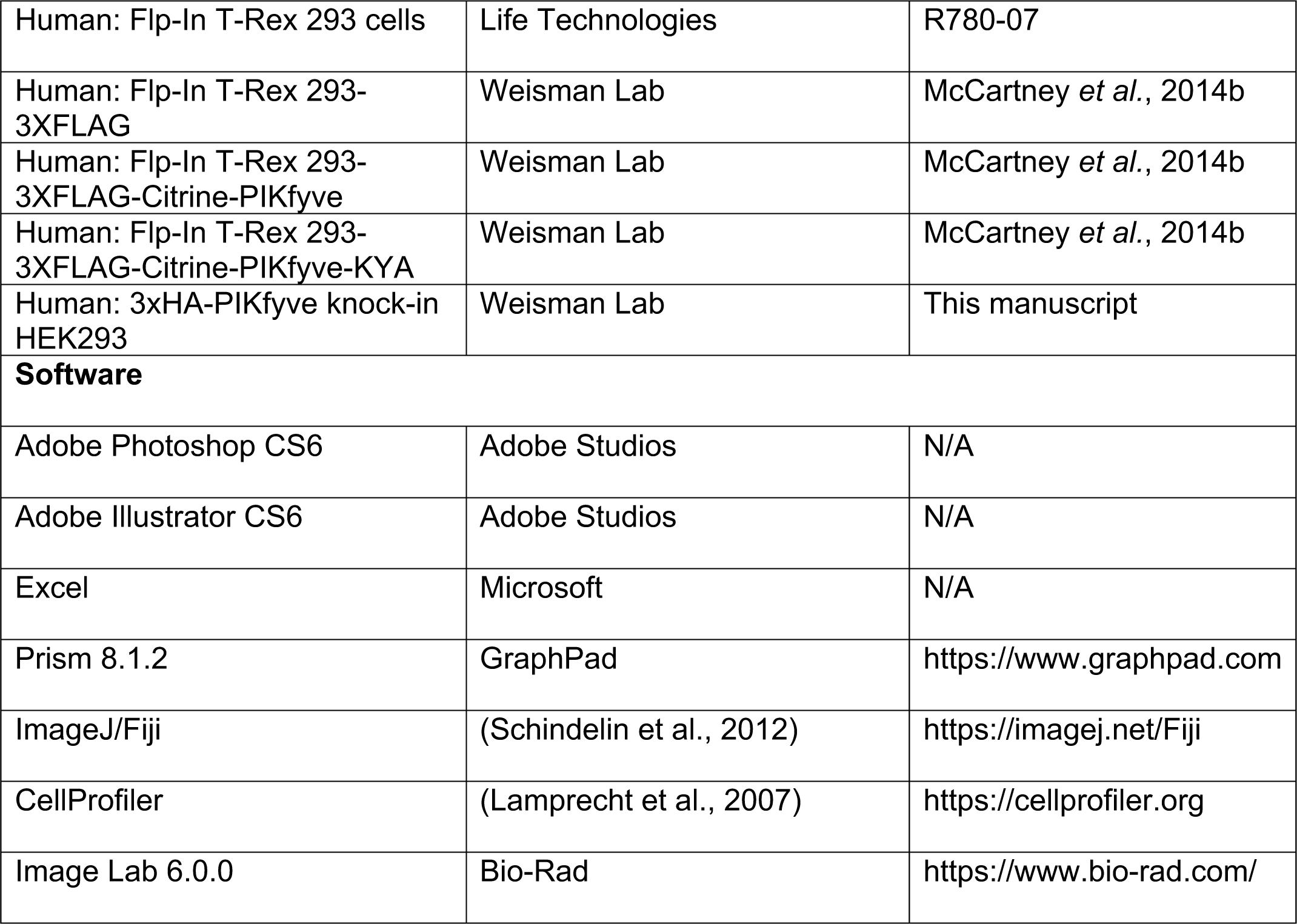

| Reagent or Resource | Company name | Catalog number |
| --- | --- | --- |
| **Antibodies** |  |  |
| Anti-HA | Cell signaling | 2367 (6E2) |
| Anti-HA | Cell signaling | 3724 (C29F4) |
| EEA1 | BD biosciences | 610456 |
| EEA1 | Cell signaling | 3288 (C45B10) |
| Lamp1 | Developmental Studies<br>Hybridoma Bank | H4A3 |
| Lamp1 | Abcam | ab24170 |
| Lamp2a | BD biosciences | 555803 (H4B4) |
| RAB4 | Abcam | ab13252 |
| RAB5 | Abcam | ab18211 |
| RAB7 | Abcam | ab137029 |
| RAB11 | Abcam | ab128913 |
| SNX17 | Protein Atlas | HPA043897 |
| FAM21 | Daniel D. Billadeau | N/A |
| CCDC93 | Proteintech | 20861-1-1AP |
| COMMD1 | R&D systems | MAB7526 |
| COMMD1 | Proteintech | 11938-1-AP |
| COMMD5 | Proteintech | 10393-1-AP |
| VPS35L | Daniel D. Billadeau | N/A |
| VPS35 | Abcam | ab10099 |
| Strumpellin | Bethyl Laboratories | A304-809A |
| β1-integrin | Millipore | MAB2000 (HB1.1) |
| β1-integrin | Santa Cruz Biotechnology | sc-13590 (P5D2) |
| β1-integrin | Cell signaling | 4706 |
| α5-integrin | Cell signaling | 4705 |
| LRP1 | Cell signaling | 64099 |
| cortactin | Millipore | 05-180 (4F11) |
| **Chemicals, Peptides, and Recombinant Proteins** |  |  |
| Apilimod | Axon Medchem | Axon1369 |
| DMEM | Thermo Fisher Scientific | 11995 |
| FBS | Sigma | F4135 |
| penicillin/streptomycin/glutamate | Life Technologies | 10378 |
| Blasticidin S HCl | Life Technologies | A11139 |
| Hygromycin B | Life Technologies | 10687 |
| Doxycycline | Sigma | D9891 |
| Fibronectin | Sigma | F1141 |
| Laminin | Gibco | 23017015 |
| **Critical Commercial Assays** |  |  |
| FuGENE 6 | Promega | E2691 |
| Lipofectamine 2000 | Thermo Fisher Scientific | 11668027 |
| Click-iT EdU Imaging Kits | Invitrogen | C10337 |
| LIVE/DEAD™<br>Viability/Cytotoxicity Kit | Invitrogen | L3224 |
| pGEM-T-Easy vector system | Promega | A1360 |
| **Recombinant DNA** |  |  |
| Plasmid: pHA-CMV | Clontech | 631604 |
| Plasmid: 6xHA-PIKfyve | Weisman Lab | This manuscript |
| Plasmid: 6xHA-PIKfyve-KYA | Weisman Lab | This manuscript |
| Plasmid: pSpCas9(BB)-2A-GFP | Addgene | 48138 |
| pLenti-Lox-EV-3XHA-COMMD1 | Weisman Lab | This manuscript |
| pLenti-Lox-EV-3XHA-COMMD1-QAA (R133Q/ H134A/ K167A) | Weisman Lab | This manuscript |
| pLenti-Lox-EV-3XHA-COMMD1-EQE (R133E/ H134Q/ K167E) | Weisman Lab | This manuscript |
| pLenti-Lox-EV-3XHA-COMMD1-EDEQE (R120E/ K129D/ R133E/ H134Q/ K167E) | Weisman Lab | This manuscript |
| **Cell lines** |  |  |
| Human: HeLa | Richard Klausner Lab |  |
| Human: HeLa | Burstein Lab | N/A |
| Human: HeLa COMMD1-/- cells | Burstein Lab | N/A |
| Human: HEK293 | ATCC | CRL-1573 |
| Human: HUH7 | David Ginsburg Lab<br>(University of Michigan-Ann Arbor) | N/A |
| Mouse: Primary MEF (Wild Type) | Weisman Lab | Zolov *et al*, 2012 |
| Mouse: Primary MEF<br>(PIKfyve<sup>β-geo/β-geo</sup>) | Weisman Lab | Zolov *et al*, 2012 |
| Mouse: Primary neonatal cardiac fibroblasts | Alan Smrcka (University of Michigan-Ann Arbor) | N/A |
| Human: Flp-In T-Rex 293 cells | Life Technologies | R780-07 |
| Human: Flp-In T-Rex 293-3XFLAG | Weisman Lab | McCartney *et al.*, 2014b |
| Human: Flp-In T-Rex 293-3XFLAG-Citrine-PIKfyve | Weisman Lab | McCartney *et al.*, 2014b |
| Human: Flp-In T-Rex 293-3XFLAG-Citrine-PIKfyve-KYA | Weisman Lab | McCartney *et al.*, 2014b |
| Human: 3xHA-PIKfyve knock-in HEK293 | Weisman Lab | This manuscript |
| **Software** |  |  |
| Adobe Photoshop CS6 | Adobe Studios | N/A |
| Adobe Illustrator CS6 | Adobe Studios | N/A |
| Excel | Microsoft | N/A |
| Prism 8.1.2 | GraphPad | https://www.graphpad.com |
| ImageJ/Fiji | (Schindelin et al., 2012) | https://imagej.net/Fiji |
| CellProfiler | (Lamprecht et al., 2007) | https://cellprofiler.org |
| Image Lab 6.0.0 | Bio-Rad | https://www.bio-rad.com/ |

### Cell culture, transfection and plasmids

HeLa, HEK293, HUH7 cell lines were grown in DMEM supplemented with 10% FBS, penicillin/streptomycin/glutamate (PSG). Primary MEF cells were cultured in DMEM containing 15% FBS and PSG. Flp-In T-Rex 293 cells that inducibly express a cDNA encoding 3xFLAG alone, 3xFLAG-Citrine-PIKfyve or 3xFLAG-Citrine-PIKfyve-KYA have been previously described (McCartney et al., 2014b). These cell lines were grown in DMEM supplemented with 10% FBS, PSG, 15 μg/mL Blasticidin S HCl and 0.4 mg/mL Hygromycin B and cells were induced with 100 ng/mL doxycycline for 12 h. For transient transfection of PIKfyve, FuGENE 6 was used following manufacture instructions and transfection time was 16-18 hours. All cells were cultured in 5% CO2 at 37°C. For immunofluorescence studies, HEK293 and 3xHA-PIKfyve knock-in HEK293 cells were grown in fibronectin and laminin-coated coverslips. 6XHA-PIKfyve and 6xHA-PIKfyve-KYA were generated by cloning PIKfyve into pHA-CMV and adding 5XHA tag using Gibson assembly as described previously(Cinato et al., 2021). HeLa COMMD1 knockout cells were generated as previously described (Singla et al., 2019) using CRISPR/Cas9-mediated gene deletion and the two COMMD1 target guides: 5’-ACGGAGCCAGCTATATCCAG-3’ AND 5’-GCGCATTCAGCAGCCCGCTC-3’.

3XHA-COMMD1, 3XHA-COMMD1-QAA, 3XHA-COMMD1-EQE and 3XHA-COMMD1-EDEQE constructs were generated by cloning cDNA of 3XHA-COMMD1 and its mutants to pLenti-Lox-EV vector using Gibson assembly. COMMD1-knockout cells were transiently transfected with these COMMD1 constructs using lipofectamine 2000 following manufacturer instructions for 24 hours.

### Generation of cells with endogenous expression of 3xHA-PIKfyve

HEK293 cells were modified by CRISPR-Cas9 genome editing to add a 3xHA tag to PIKfyve at the N-terminus. Donor DNA spanning 315 base pairs on the left homologous region and 353 base pairs on the right homologous region was generated by overlap extension PCR and then cloned into the pGEM-T-Easy vector system. Guide RNA (TGATAAGACGTCCCCAACAC) for PIKfyve was cloned into pX458, a pSpCas9-2A-EGFP vector. Lipofectamine 2000 was used for transfection of cells with pX458 expressing Cas9 along with a gRNA donor vector. Three days after transfection GFP-positive single cells were sorted using flow cytometry into a 96-well plate containing conditioned media. 3xHA-PIKfyve knock-in HEK293 cells were validated by PCR, the PCR product was sequence verified and the cell lysate were verified by western blot.

### Wound healing and cell spreading assays

Cells were grown on coverslips or fibronectin-coated plastic dishes to full confluency and were wounded using a pipet tip, then incubated in DMEM with 10% FBS for the indicated treatments and time points. Quantification of wound healing was performed on 5 random fields for each condition and wound area closure was quantified from three independent experiments. For cell spreading assays, cells were trypsinized, seeded onto fibronectin-coated plastic dishes and allowed to attach to dishes in DMEM with 10% FBS for the indicated times.

### Cell proliferation and cell viability assay

The cell proliferation assay was performed with Click-iT EdU Imaging Kits. HeLa cells grown on coverslips with treatments as indicated were incubated with 10 µM EdU (5-ethynyl-2’-deoxyuridine) in DMEM with 10% FBS for 30 min. Cells were fixed, permeabilized and incubated with Click-iT reaction cocktail to detect the incorporated EdU according to manufacture instructions. Cells were mounted and analyzed. Cells which incorporated Edu were identified as proliferating.

The cell viability assay was performed using LIVE/DEAD™ Viability/Cytotoxicity Kit, for mammalian cells. HeLa cells grown on coverslips with treatments as indicated were incubated in DMEM with 10% FBS containing 2 µM calcein AM and 4 µM ethidium homodimer-1 (EthD-1) for 30 min. Cells were mounted with 10 µl PBS and analyzed by fluorescence microscopy.

### Surface biotinylation

Cells were incubated with DMSO or 1 µM apilimod for 1 h at 37^0^C, then transferred to 4^0^C for 15 min. Cells were washed with ice cold wash buffer (PBS containing 2.5 mM MgCl_2_ and 1mM CaCl_2_) and incubated with ice cold 0.5 µg/ml NHS-SS-Biotin (Pierce) for 20 min. Biotinylation was quenched by incubating the cells with ice cold 100 mM glycine for 10 min. Cells were then pelleted, lysed in RIPA buffer (Pierce) containing protease and phosphatase inhibitors. 3 mg of protein lysate was incubated overnight with 100 µl of streptavidin bead slurry. Western blot analysis was performed on 20% of the total of each immunoprecipitate, and 50 µg of each lysate.

### Labeling of surface exposed integrin

HeLa cells after indicated treatments were incubated with ice cold serum free media containing 0.5% BSA and 5 µg/ml mouse anti-β1-integrin antibody (MAB2000) for 1 h at 4^0^C. Cells were fixed in ice cold 4% paraformaldehyde for 30 min at 4^0^C. Cells were permeabilized, and immunostained with Alex Fluor 488-conjugated donkey anti-mouse secondary antibodies. To determine the effect of PIKfyve mutants on the surface levels of integrin, cells were fed with complete media for 2 hours prior to labelling surface integrin. Cells were then incubated with 5 µg/ml mouse anti-β1-integrin antibody for 1 h at 4^0^C. Cells were fixed at 4^0^C for 30 min and permeabilized. Cells were immunostained with rabbit anti-HA antibodies followed by Alex Fluor 488-conjugated donkey anti-mouse and Alex Fluor 488-conjugated goat anti-rabbit secondary antibodies.

### Integrin trafficking experiments

To determine the dynamic regulation of integrin levels at the cell surface, cells were incubated with serum free media containing 0.5% BSA and 5 µg/ml mouse anti-β1-integrin antibody (MAB2000) for 1 h at 4^0^C. Cells were then incubated with DMSO or 1 µM apilimod in DMEM with 10% FBS at 37^0^C for the indicated times. To measure the surface levels, cells were fixed with 4% paraformaldehyde after treatment. To determine the internalized and unrecycled pool, cells were fixed after a brief acid wash of 0.5% acetic acid and 0.5 M NaCl for 1 min, to remove surface antibodies. Fixed cells were immunostained with secondary antibodies and analyzed by confocal microscope.

To measure β1-integrin recycling, cells were incubated with 5 µg/ml mouse anti-integrin antibody (P5D2) in DMEM with 10% FBS for 1 h at 37^0^C. Cells were acid washed with PBS, pH 3.0 for 1 min. At this stage, cells were either fixed with 4% paraformaldehyde to determine the internalize pool of β1-integrin or the following treatments were performed. Cells were then either untreated or incubated with DMSO or 1 µM apilimod in DMEM with 10% FBS at 37^0^C for the indicated times. To determine the non-recycled pool, cells were fixed after a brief acid wash with PBS, pH 3.0 for 1 min. Fixed cells were incubated with the indicated antibodies and analyzed. Cells were co-labeled with Texas Red-WGA to mark the cell border. To determine the localization of non-recycled integrin, cells were co-incubated with antibodies to the endocytic markers: EEA1 for early endosome, LAMP1 for late endosome and lysosome, RAB4 for fast recycling endosomes and RAB11 for slow recycling endosomes.

### Immunofluorescence, and image acquisition

For PIKfyve colocalization studies, cells were serum starved for 2 hours and fed with DMEM with 10% FBS for 30 min prior to fixation; other cells were fixed directly after the indicated treatments. Cells were fixed with 4% paraformaldehyde for 10 min at room temperature unless specified. Fixed cells were permeabilized with PBS containing 0.5% BSA and 0.2% saponin. Permeabilized cells were incubated with the indicated antibodies and analyzed. Images were acquired with an Olympus FV1000, LEICA SP5 or LEICA SP8 confocal microscope under an oil immersion 60x or 63X objective, respectively. Immunoblots were imaged using a ChemiDoc Imaging System (Bio-Rad)

### Image analysis

Images were analyzed using Fiji (ImageJ; NIH) or CellProfiler. To determine the intensity of integrin during trafficking and surface labeling experiments, cells were segmented using the ImageJ crop function and integrated density was measured using the analyze function. The changes in the surface levels of integrin were inferred from the percentage of integrin intensity within 0.8 μm from the plasma membrane; the enlarge function was employed to mark this area. To measure the intensity of proteins in VPS35-positive endosomes, masks were created in ImageJ using VPS35 fluorescence and overlaid onto the fluorescence channel of the protein of interest. To determine the colocalization between integrin and endocytic markers, images were segmented, and colocalization was measured using CellProfiler. To determine the colocalization of PIKfyve with other proteins or markers, images were cropped and the colocalization was measured using the Jacop plugin-in, ImageJ. Immunoblots were analyzed using Image Lab Software.

### Statistical analysis

All experiments were performed at least three times. Statistical analyses are described in the figure legends. Statistical analyses were performed in GraphPad Prism 8.1.2.

