## Supplemental Figures for "Lipid kinases VPS34 and PIKfyve coordinate a phosphoinositide cascade to regulate Retriever-mediated recycling on endosomes"

### Figure S1, related to Figure 1

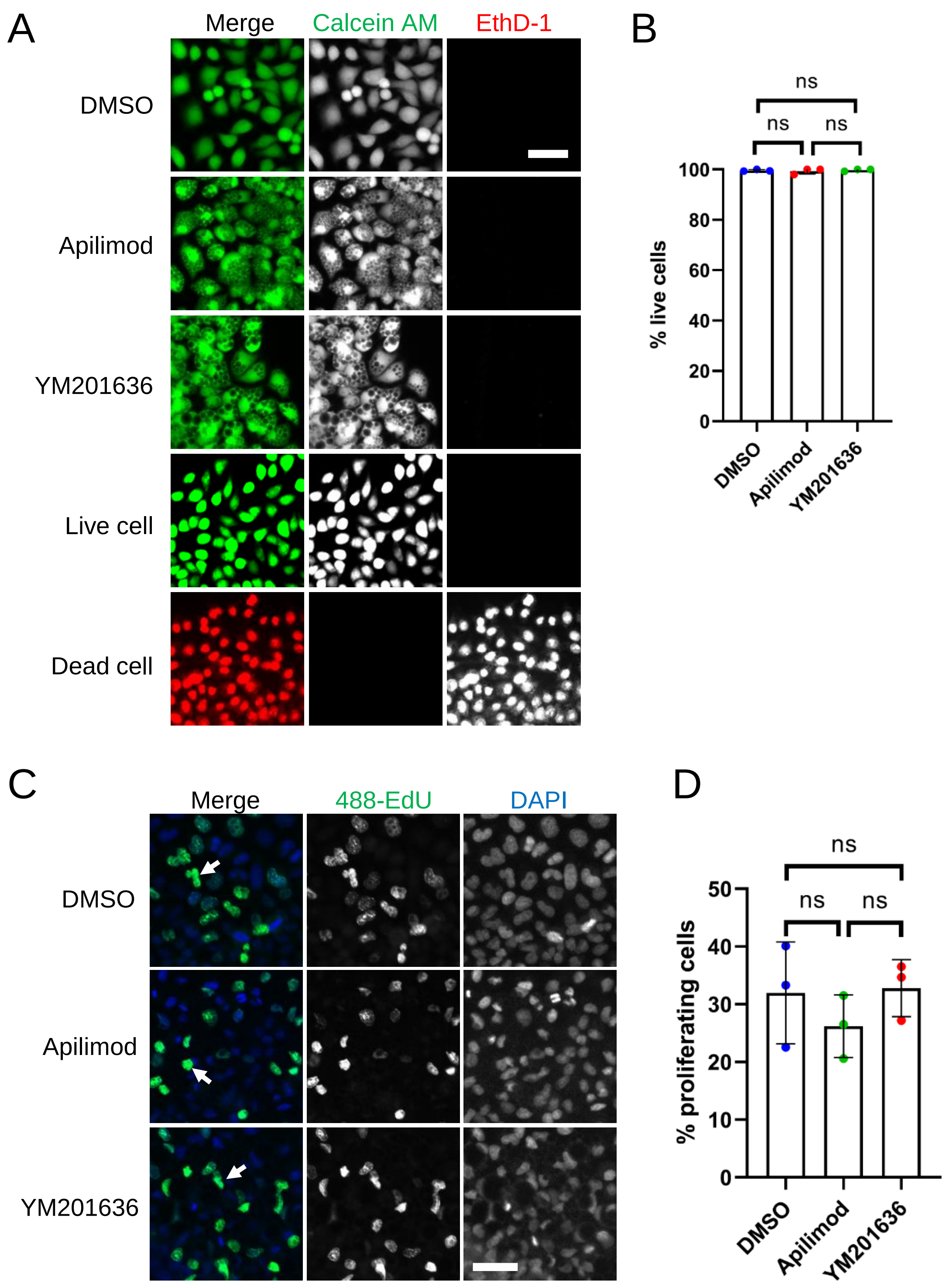

**Figure S1. Inhibition of PIKfyve does not affect cell viability or proliferation.**

A-B. Cell viability was assessed for HeLa cells that were either treated with DMSO, 1 µM apilimod or 0.8 µM YM201636 for 27 h. Untreated cells and cells treated with methanol for 20 min were used as live and dead cell controls, respectively. Percentage of live cells was quantified.

C. Cell proliferation was measured in HeLa cells treated with DMSO, 1 µM apilimod or 0.8 µM YM201636 for 27 h.

D. The percentage of proliferating cells in (C) was quantified.

Data presented as the mean ± SE. Statistical significance from three independent experiments was analyzed using one-way ANOVA and Tukey post hoc tests (B, D). ns, not significant. Bar-100 µm.

### Figure S2, related to Figure 1

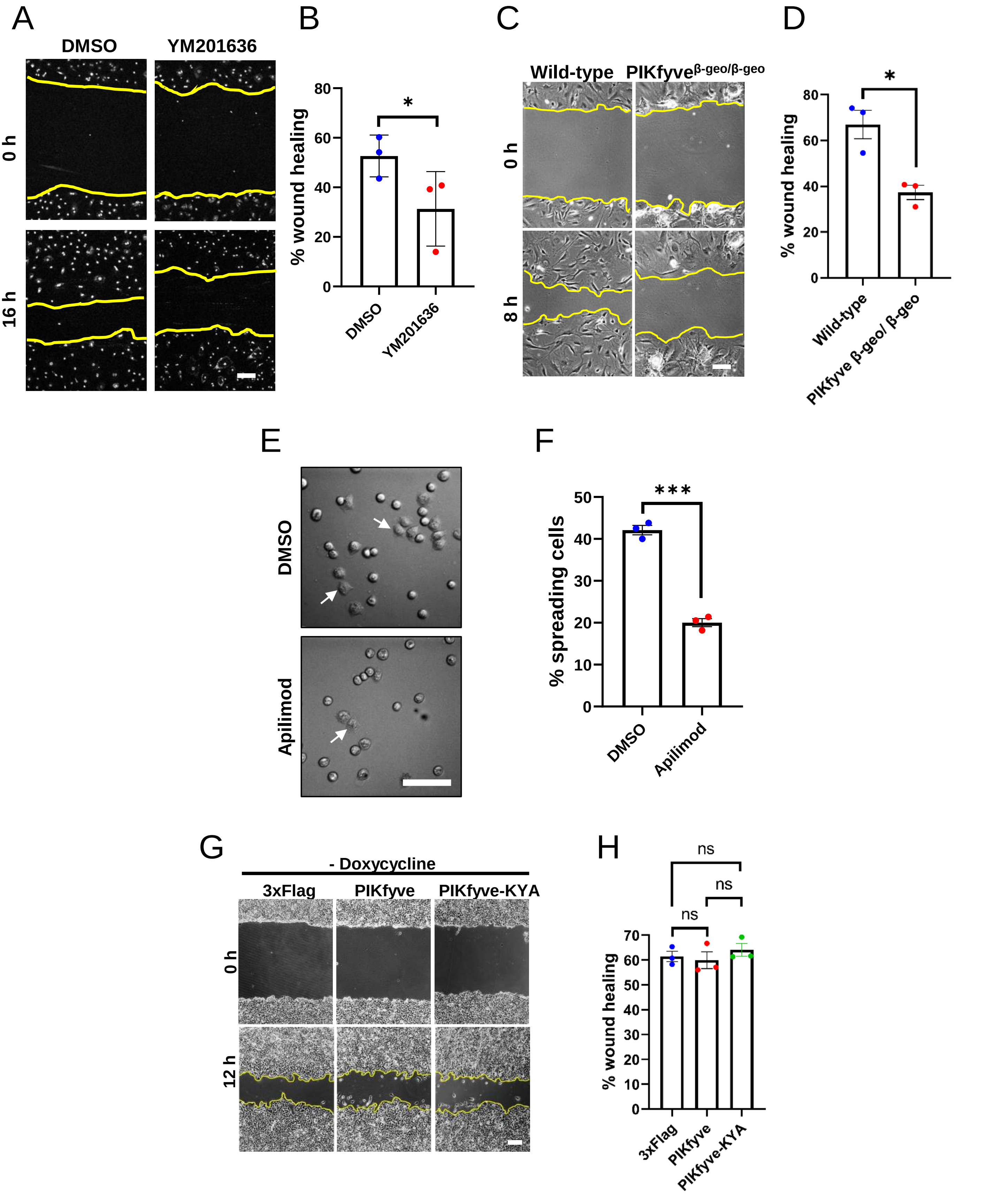

**Figure S2. PIKfyve is required for cell migration and cell adhesion.**

A-D. PIKfyve is required for cell migration. (A) Wound healing assays were performed on primary neonatal cardiac fibroblasts in the presence of DMSO or 0.8 µM YM201636 for 16 h. (B) Percentage of wound closure was quantified. (C) Wound healing assays were performed for 8 h on primary MEF cells derived from wild-type and hypomorphic PIKfyve$^{\beta\text{-geo}/\beta\text{-geo}}$ mice. (D) Percentage of wound area closure was quantified.

E-F. PIKfyve is required for cell spreading. (E) HeLa cells were trypsinized, seeded in media containing either DMSO or 1 µM apilimod and incubated for 1 h. Arrows indicate examples of cells that exhibited spreading. (F) Percentage of cells that spread onto the surface were quantified.

G-H. Cell migration is not elevated when there is no induction of the hyperactive mutant of PIKfyve**.** Wound healing assay was performed on HEK293T cells stably expressing doxycycline-inducible wild-type PIKfyve or hyper-active PIKfyve-KYA in the absence of doxycycline for 12 h. Percentage of wound area closure was quantified. Yellow lines indicate the migration front.

Data presented as mean ± SE. Statistical significance from three independent experiments was determined using paired two-tailed Student's T-Test (B, D, F) or one-way ANOVA and Tukey post hoc tests (H). *** $P<0.005$, * $P<0.05$, and ns, not significant. Bar-100 µm. Error Bar, SE.

### Figure S3, related to Figure 3 and 6

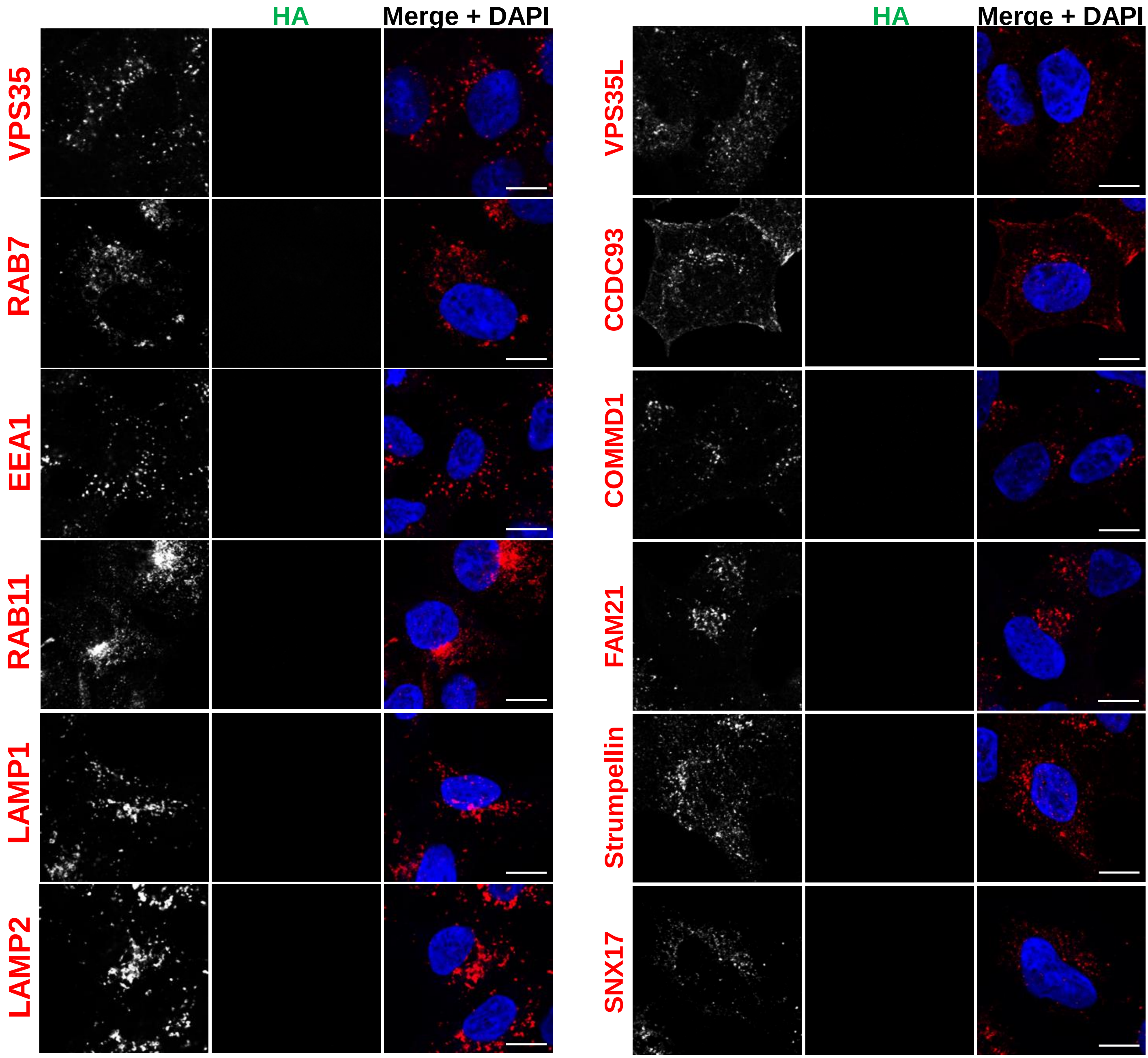


**Figure S3. Immunofluorescence localization of endosomal proteins in unedited HEK293 cells (control for Figure 3 and Figure 6).** HEK293 cells were fixed, permeabilized and incubated with antibodies against HA and against markers for either the retromer (VPS35), early endosomes (EEA1), late endosomes (RAB7), recycling endosomes (RAB11), lysosomes (LAMP1 and LAMP2), SNX17, the Retriever subunit, VPS35L, the CCC complex subunits, COMMD1 and CCDC93, or the WASH complex subunits, Strumpellin and FAM21.

### Figure S4, related to Figure 5

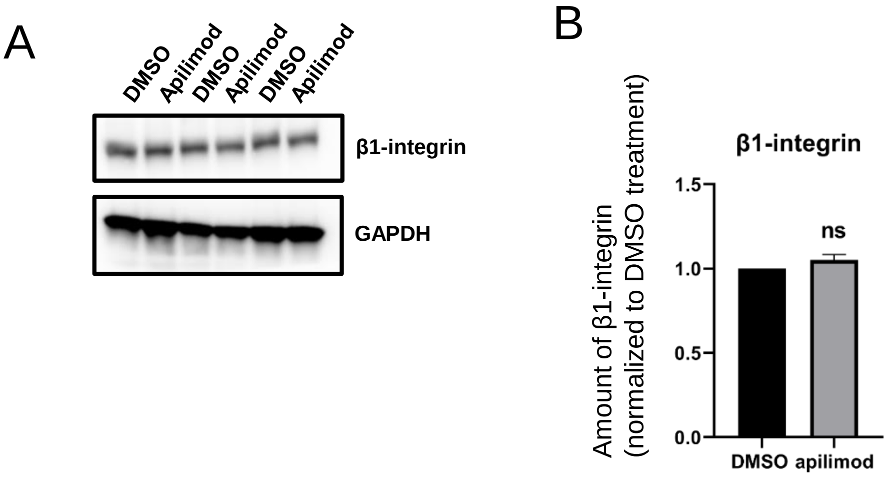


**Figure S4.** β**1-integrin levels remain stable during PIKfyve inhibition.**

A-B. HeLa cells were treated with DMSO or 1 µM apilimod for 30 min. (A) Lysates from three independent experiments were immunoblotted with antibodies against β1-integrin and GAPDH. (B) Levels of β1-integrin were quantified and values were normalized to DMSO control.

Data presented as mean ± SE. Statistical significance was analyzed using paired two-tailed Student's T-Test. ns, not significant.

### Figure S5, related to Figure 7

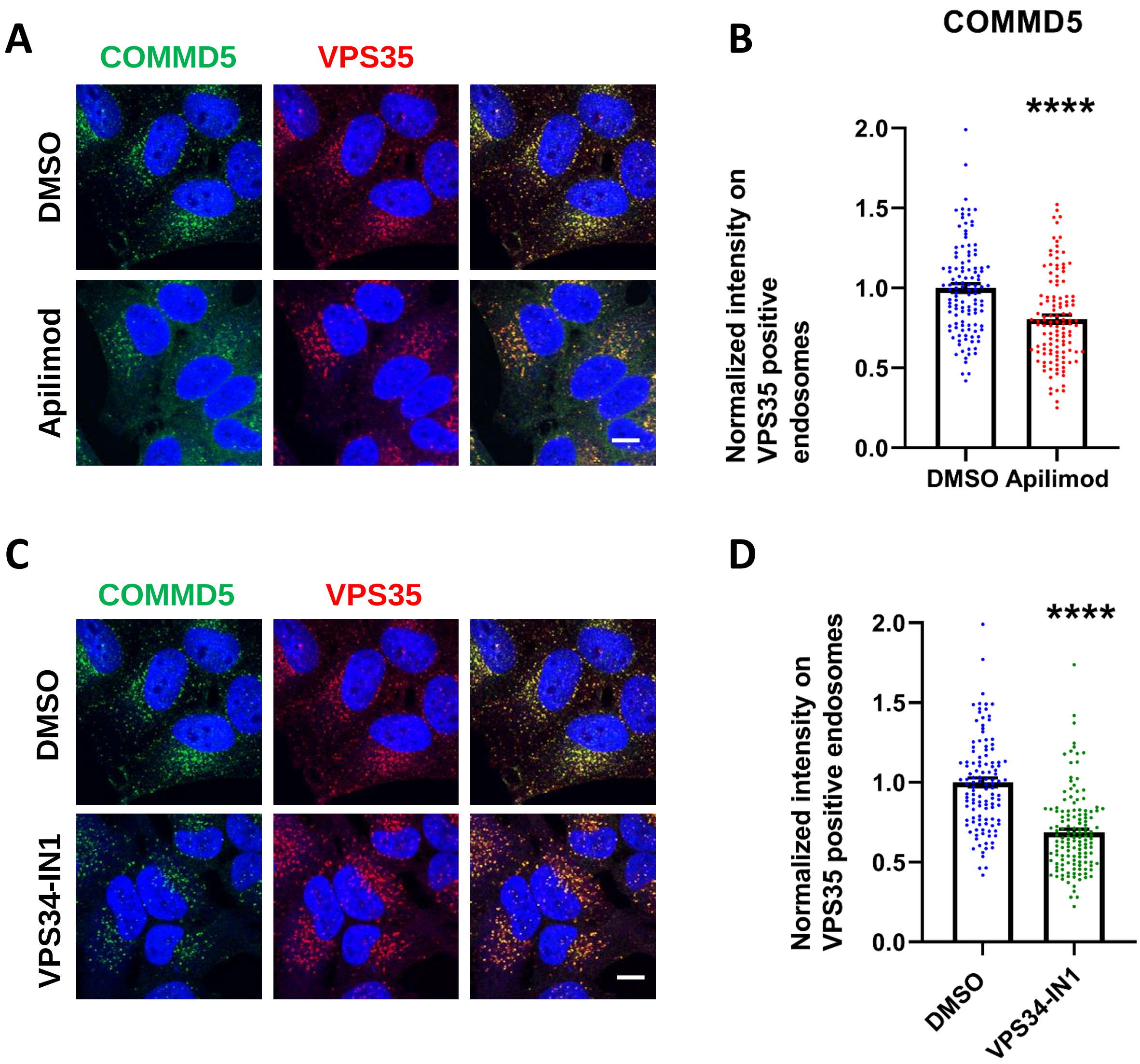


**Figure S5. Inhibition of VPS34 or PIKfyve results in a loss of the CCC complex subunit, COMMD5 from endosomes.**

HeLa cells treated with DMSO , 1 µM apilimod (A-B) or 1 µM VPS34-IN1 (C-D) for 30 min were fixed, permeabilized and co-stained with antibodies against VPS35 and COMMD5. The intensity of COMMD5 on VPS35-positive endosomes was quantified and values were normalized to the corresponding average intensity of the DMSO treatment cohort (B and D).

Data presented as mean ± SE. Statistical significance from three independent experiments were analyzed using unpaired two-tailed Student's T-test. *** $P<0.005$ and **** $P<0.001$, and ns not significant. Bar-10 µm.

### Figure S6, related to Figure 7

A

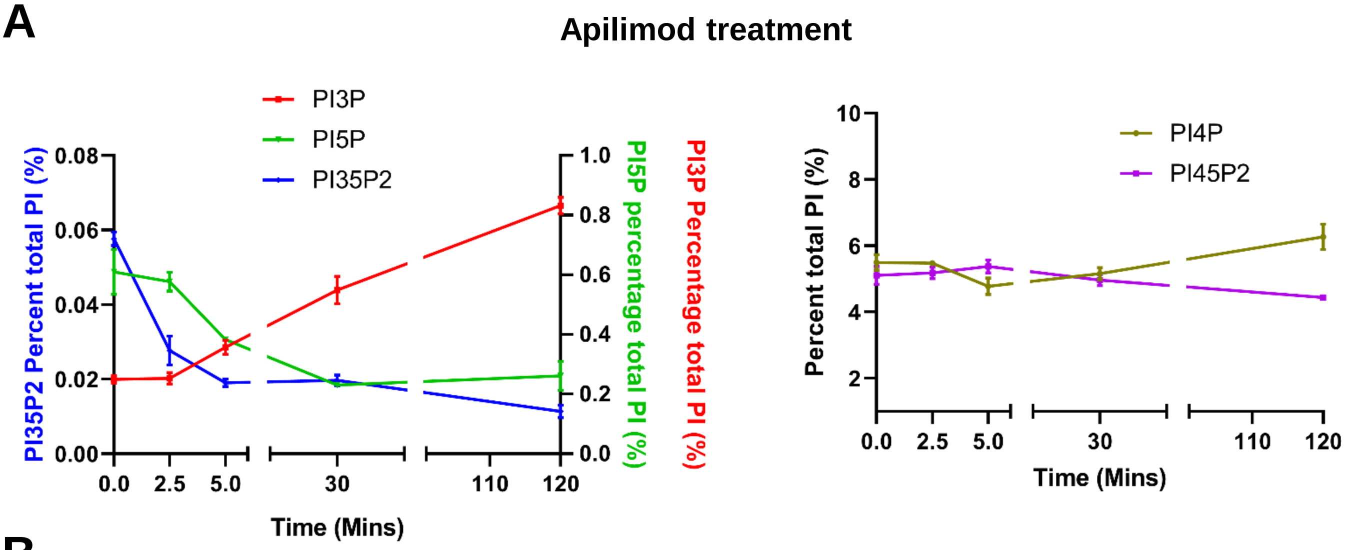


B

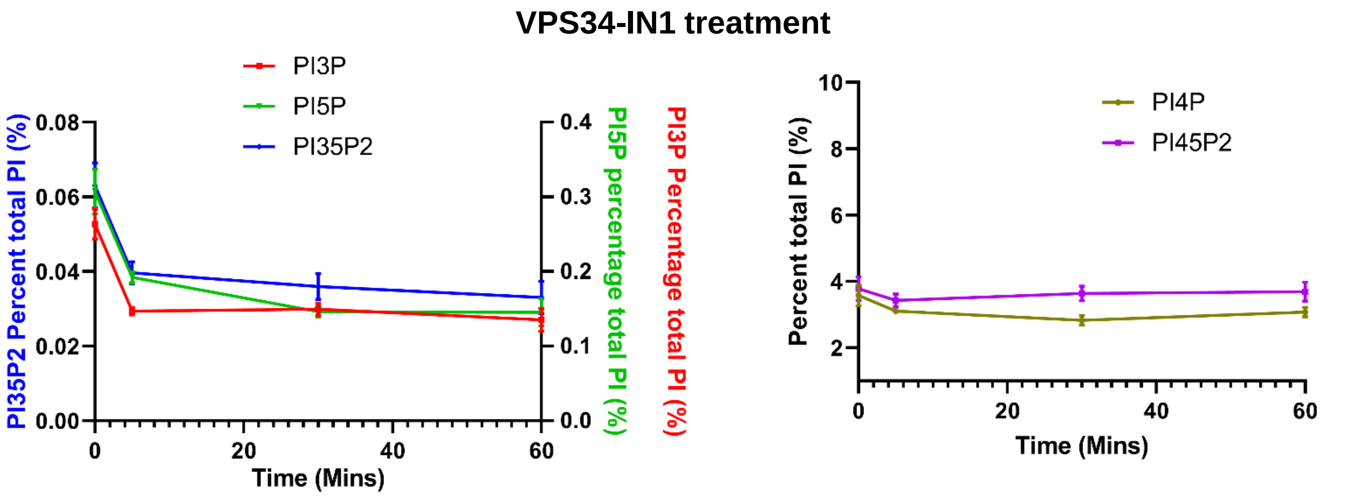


**Figure S6. Apilimod and VPS34-IN1 are potent inhibitors of PIKfyve and VPS34 respectively.**

A. MEFs cells were incubated with myo-[2-$H^3$] inositol labelled media for 48 hours and cells were either untreated or treated with 1 µM apilimod for the last 2.5, 5, 30 or 120 minutes of the labeling. Note that PI3P is elevated approximately 4-fold at 120 min of treatment. Levels of individual phosphoinositide lipids were quantified (data adapted from McCartney et al., 2014).

B. MEFs cells were incubated with myo-[2-$H^3$] inositol labelled media for 48 hours and cells were either untreated or treated wiith 1 µM VPS34-IN1 for the last 5, 30 or 60 minutes of the labeling. Levels of individual phosphoinositide lipids were quantified.

Data presented as mean ± SE from three independent experiments.

### Figure S7 related to Figure 7

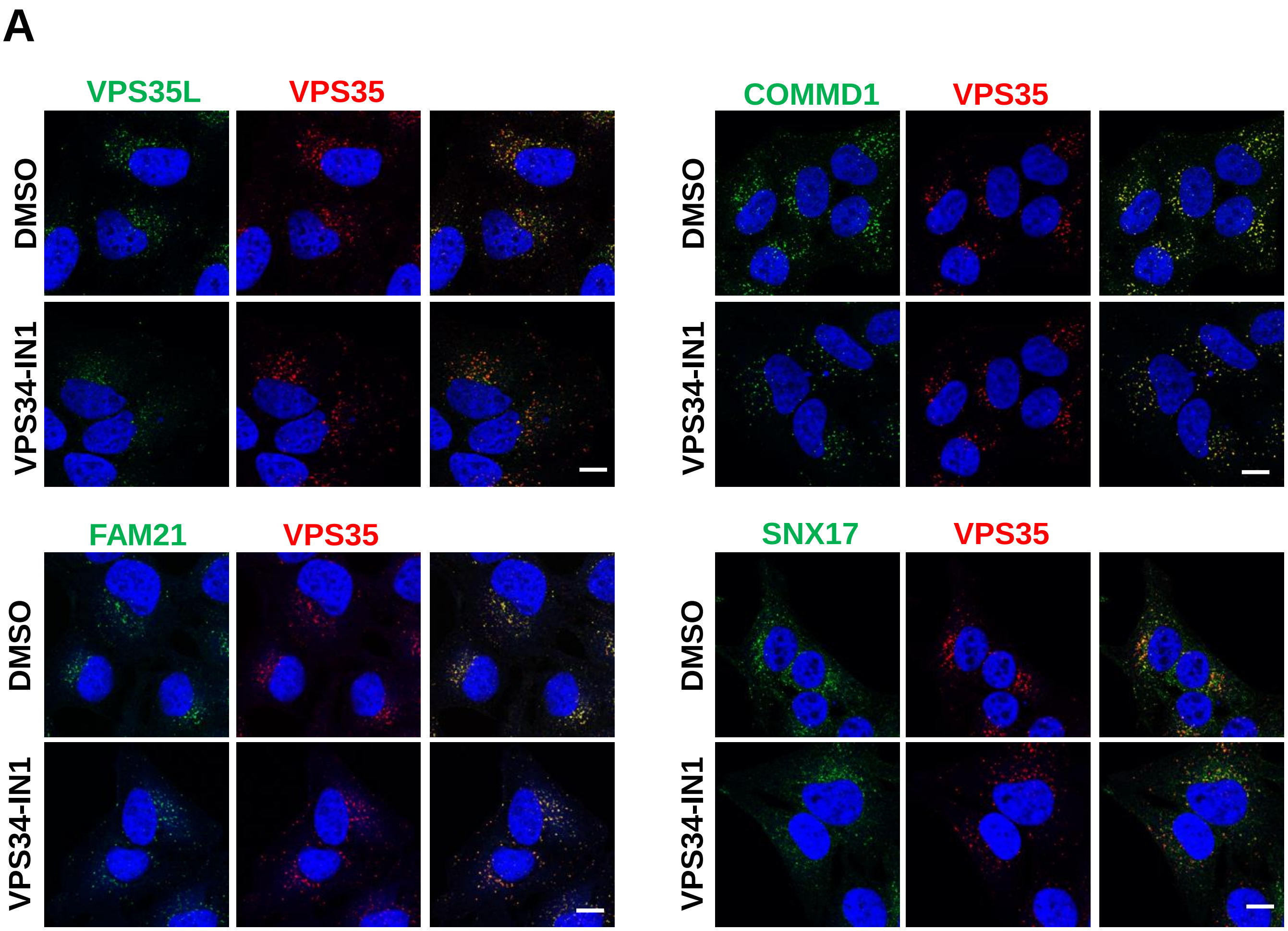


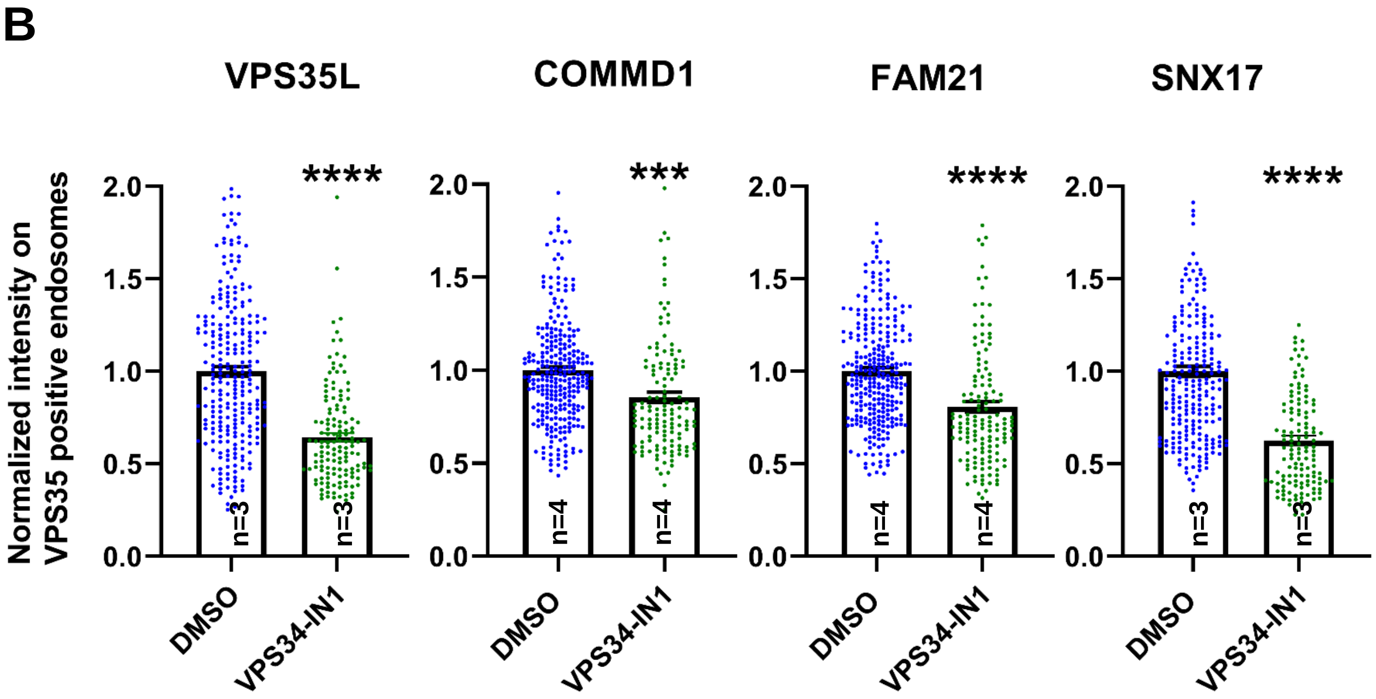

**Figure S7. Inhibition of the PI3-kinase, VPS34 results in a loss of SNX17, CCC, Retriever and WASH complexes from endosomes.**

A. HeLa cells treated with DMSO or 1 µM VPS34-IN1 for 30 min were fixed, permeabilized and co-stained with antibodies against VPS35 (A-D) and antibodies against either COMMD1, VPS35L, FAM21 and SNX17.

B. The intensity of COMMD1, VPS35L, FAM21 and SNX17 on VPS35-positive endosomes was quantified and values were normalized to the corresponding average intensity of the DMSO treatment cohort.

Data presented as mean ± SE. Statistical significance from three or four independent experiments were analyzed using unpaired two-tailed Student's T-test. *** P<0.005 and **** P<0.001, and ns not significant. Bar-10 µm.

### Figure S8, related to Figure 7

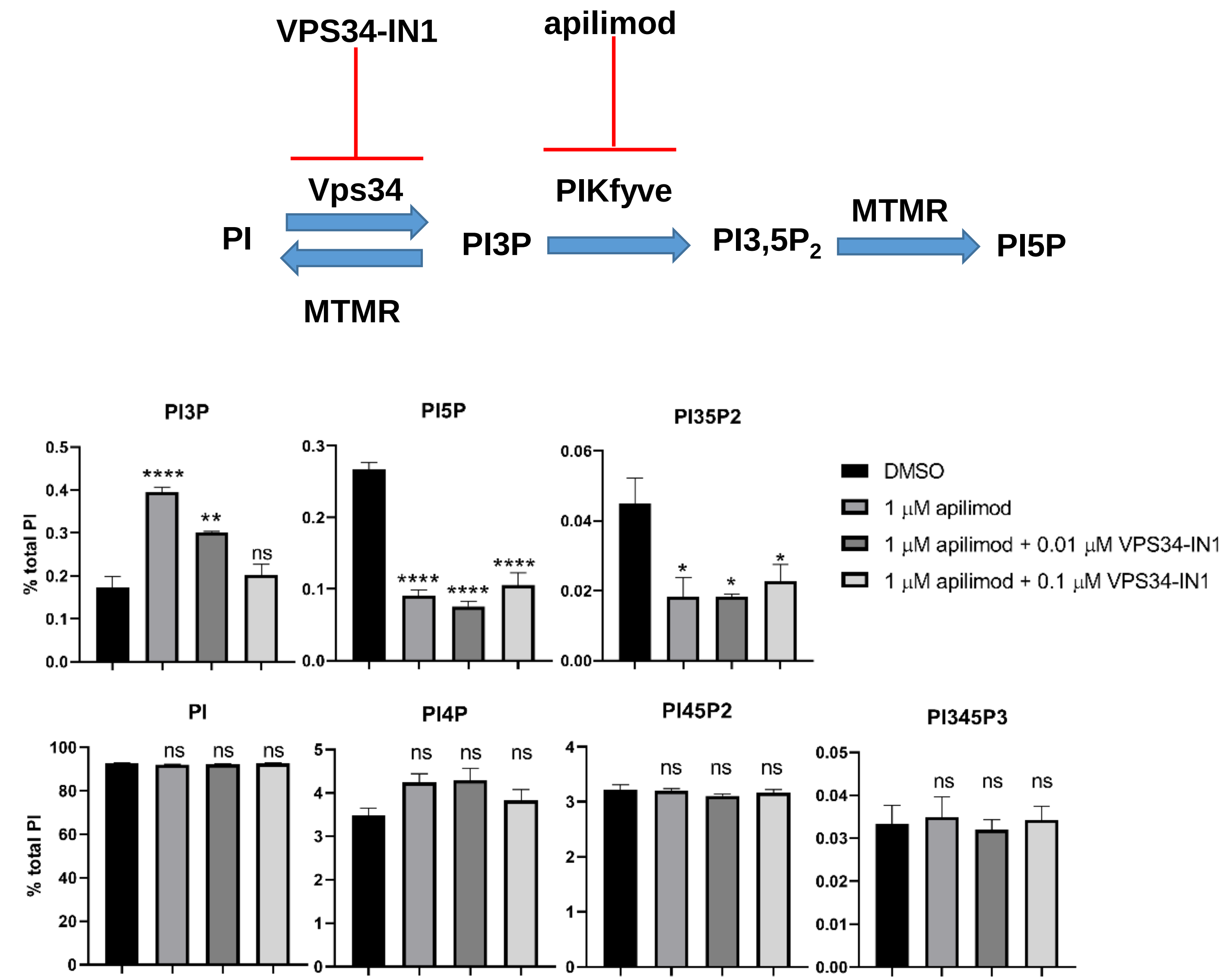


**Figure S8. Partial inhibition of VPS34 with 0.1 μM VPS34-IN1 combined with treatment with apilimod prevents the elevation of total cellular pools of PI3P.**

MEFs cells were incubated with myo-[2-$H^3$] inositol labelled media for 48 hours. Cells were either treated with DMSO, 1 μM apilimod or co-treated with 1 μM apilimod and 0.1 μM or 0.01 μM VPS34-IN1 for the last 30 minutes of the labeling. Levels of individual phosphoinositide lipids were quantified.

Data presented as mean ± SE. Statistical significance from three independent experiments were analyzed using one-way ANOVA and Tukey post hoc tests. * P<0.05, ** P<0.01, *** P<0.005 and **** P<0.001, and ns not significant.

### Figure S9, related to Figure 8

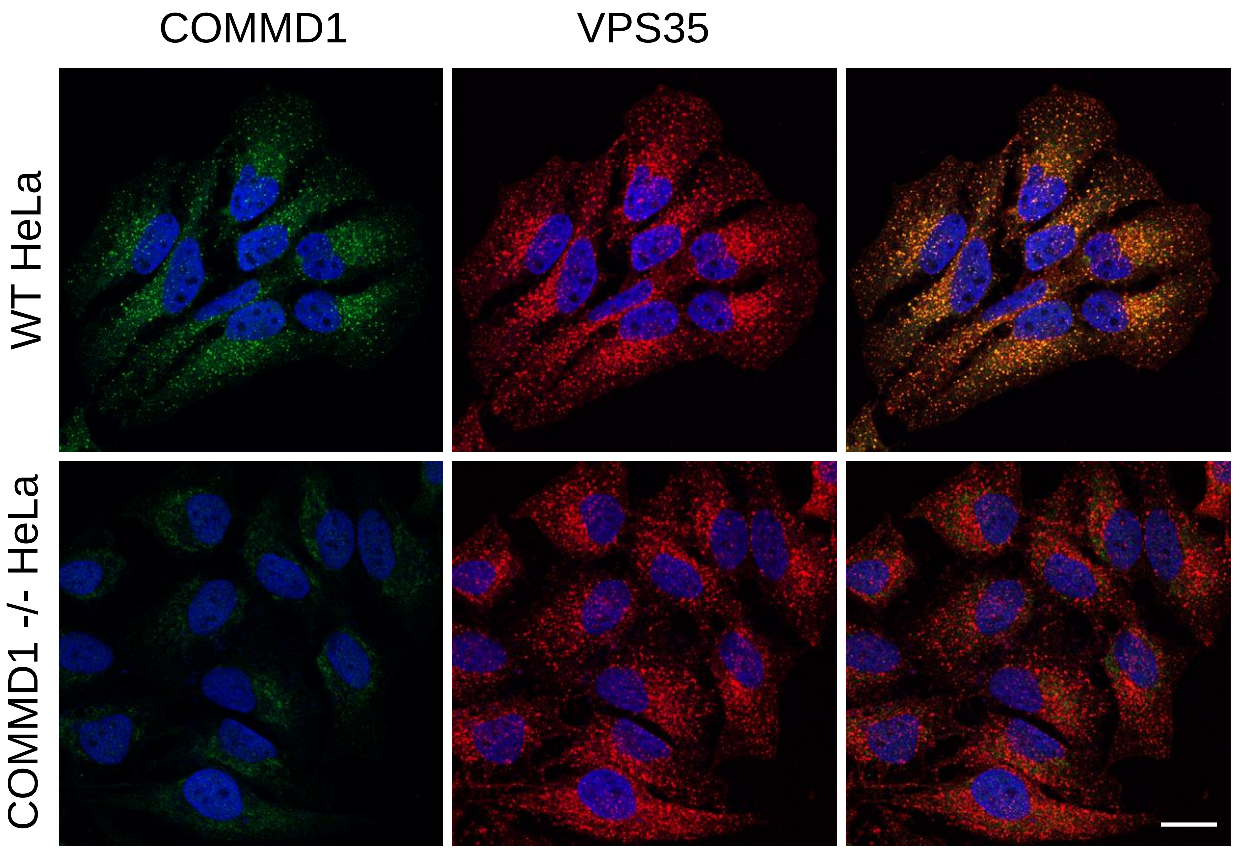


**Figure S9. Validation of COMMD1-/-  knock-out HeLa cells.**

Wild type and COMMD1-/- HeLa cells were fixed, permeabilized and immunostained with antibodies against COMMD1 and VPS35. Bar 20 µm

### Figure S10

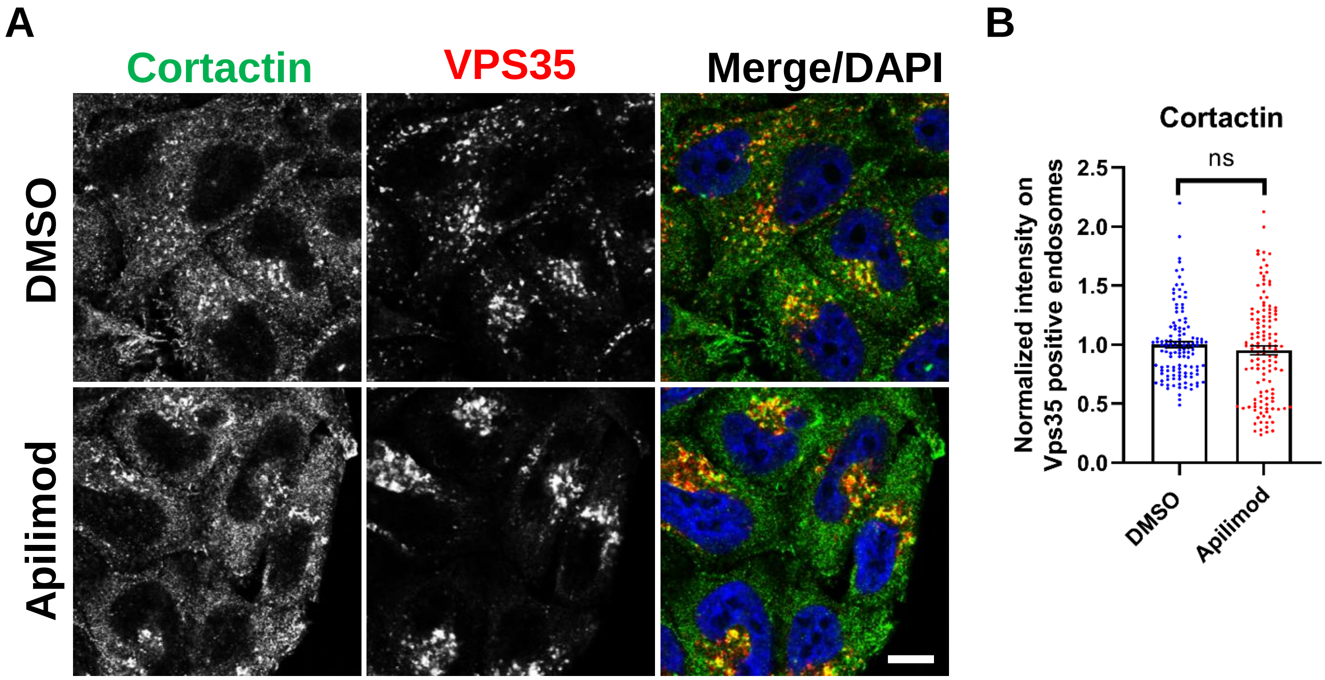


**Figure S10. Acute inhibition of PIKfyve does not alter cortactin co-localization on Vps35 endosomes.**

A. HeLa cells grown on coverslips were treated with either DMSO or 1 µM apilimod for 30 min. Cells were fixed, permeabilized and incubated with antibodies against cortactin and VPS35. Scale bar, 10 µm.

B. The intensity of cortactin on VPS35-positive endosomes was quantified and values were normalized to the average of DMSO-treated cells.

Data presented as mean ± SE. Statistical significance of data from three independent experiments was analyzed using an unpaired two-tailed Student’s T-Test. ns, not significant.
